# Motor planning modulates neural activity patterns in early human auditory cortex

**DOI:** 10.1101/682609

**Authors:** Daniel J. Gale, Corson N. Areshenkoff, Claire Honda, Ingrid S. Johnsrude, J. Randall Flanagan, Jason P. Gallivan

**Author notes:** Correspondence should be addressed to: Jason Gallivan, Centre for Neuroscience Studies, Queen’s University.

## Abstract

It is well established that movement planning recruits motor-related cortical brain areas in preparation for the forthcoming action. Given that an integral component to the control of action is the processing of sensory information throughout movement, we predicted that movement planning might also modulate early sensory cortical areas, readying them for sensory processing during the unfolding action. To test this hypothesis, we performed two human functional MRI studies involving separate delayed movement tasks and focused on pre-movement neural activity in early auditory cortex, given its direct connections to the motor system and evidence that it is modulated by motor cortex during movement in rodents. We show that effector-specific information (i.e., movements of the left vs. right hand in Experiment 1, and movements of the hand vs. eye in Experiment 2) can be decoded, well before movement, from neural activity in early auditory cortex. We find that this motor-related information is represented in a separate subregion of auditory cortex than sensory-related information and is present even when movements are cued visually instead of auditorily. These findings suggest that action planning, in addition to preparing the motor system for movement, involves selectively modulating primary sensory areas based on the intended action.

## INTRODUCTION

Most theories of motor control distinguish between the planning of a movement and its subsequent execution. Research examining the neural basis of movement planning has commonly used delayed movement tasks—in which instructions about what movement to perform are separated in time from the instruction to initiate that movement—and has focused on delay period activity in motor-related brain areas. The main focus of this work has been on understanding how planning-related neural activity relates to features of the forthcoming movement to be executed (e.g., Tanji and Evarts 1976; Messier and Kalaska 2000a), and more recently, how the dynamics of these neural activity patterns prepare the motor system for movement (Churchland et al. 2012). However, in addition to generating appropriate muscle commands, a critical component of skilled action is the prediction of the sensory consequences of movement (Wolpert and Flanagan 2001), which is thought to rely on internal models (Wolpert and Miall 1996). For example, the sensorimotor control of object manipulation tasks involves predicting the sensory signals associated with contact events, which define subgoals of the task (Flanagan et al. 2006). Importantly, these signals can occur in multiple sensory modalities, including tactile, proprioceptive, visual, and auditory (Johansson and Flanagan 2009). By comparing the predicted to actual sensory outcomes, the brain can monitor task progression, detect performance errors, and quickly launch appropriate, task-protective corrective actions as needed (Johansson and Flanagan 2009). For instance, when lifting an object that is heavier than expected, anticipated tactile events, associated with lift-off, fail to occur at the expected time, triggering a corrective response. Likewise, similar compensatory behaviour has also been shown to occur during action-related tasks when anticipated auditory events fail to occur at the predicted time (Safstrom et al. 2014). Sensory prediction is also critical in sensory cancellation, such as the attenuation of predictable sensory events that arise as a consequence of movement. Such attenuation is thought to allow the brain to disambiguate sensory events due to movement from events due to external sources (Schneider and Mooney 2018a).

At the neural level, anticipation of the sensory consequences of movement has been shown to rely on a corollary discharge signal or efference copy of outgoing motor commands being sent, in parallel, to early sensory systems (Holst et al. 1950; Crapse and Sommer 2008). Consistent with this idea, work in crickets, rodents, songbirds and nonhuman primates has reported modulations in neural activity in early sensory areas such as auditory (Poulet and Hedwig 2002; Eliades and Wang 2008; Mandelblat-Cerf et al. 2014; Schneider et al. 2014) and visual cortex (Saleem et al. 2013; Lee et al. 2014; Leinweber et al. 2017) during self-generated movements. Critically, these movement-dependent modulations are distinct from modulations related to sensory reafference and have even been shown to occur when sensory reafference is either masked (Schneider et al. 2014; Leinweber et al. 2017) or removed entirely (Keller et al. 2012; Keck et al. 2013; Saleem et al. 2013). This importantly demonstrates that these modulations result from automatic mechanisms that are motor in origin (Leinweber et al. 2017; Schneider and Mooney 2018a). Given the functional importance of predicting task-specific sensory consequences, we hypothesized that action planning, in addition to preparing motor areas for execution (Churchland et al. 2010; Shenoy et al. 2013a), involves the automatic preparation of primary sensory areas for processing task-specific sensory signals. Given that these sensory signals will generally depend on the precise action being performed, this hypothesis predicts that neural activity in early sensory areas will represent specific motor-related information prior to the movement being executed, during action preparation.

One sensory system that is particularly well suited to directly testing this hypothesis is the auditory cortex. The mammalian auditory system exhibits an extensive, highly interconnected web of feedback projections, providing it with access to the output of information processing across multiple distributed brain areas (Hackett 2015). To date, this feedback architecture has been mainly implicated in supporting auditory attention and working memory processes (Linke and Cusack 2015; Kumar et al. 2016). However, recent work in rodents has also demonstrated that the auditory cortex receives significant direct projections from ipsilateral motor cortex (Nelson et al. 2013a; Schneider et al. 2014, 2018). Consistent with this coupling between the motor and auditory systems, recent studies in both humans and rodents have shown that auditory cortex is functionally modulated by top-down motor inputs during movement execution (Reznik et al. 2014, 2015; Schneider et al. 2014, 2018). While the focus of this prior work has been on the real-time attenuation, during movement execution, of the predictable auditory consequences of movement, it did not selectively focus on the movement planning process itself, or the broader function of the motor system in modulating early auditory activity in preparation for action.

Here we show, using functional MRI and two separate delayed movement experiments involving object manipulation, that the movement effector to be used in an upcoming action can be decoded from delay period activity in early auditory cortex. Critically, this delay period decoding occurred despite the fact that any auditory consequences associated with movement execution (i.e., the sounds associated with object lifting and replacement) were completely masked both by the loud MRI scanner noise and the headphones that participants wore to protect their hearing. This indicates that the pre-movement decoding in the auditory cortex is automatic in nature (i.e., occurs in the absence of any sensory reafference) and of motor origin. Beyond real-time sensory attenuation, these findings suggest that, during movement preparation, the motor system selectively changes the neural state of early auditory cortex in accordance with the specific motor action being prepared, likely readying it for the processing of sensory information that normally arises during subsequent movement execution.

## MATERIALS & METHODS

### Overview

To test our hypothesis that motor planning modulates the neural state of early auditory cortex, we performed two separate functional MRI experiments that used delayed movement tasks. This allowed us to separate motor planning-related modulations from the later motor and somatosensory-related modulations that occur during movement execution. In effect, these delayed movement tasks allowed us to ask whether the motor command being prepared—but not yet executed—can be decoded from neural activity patterns in early auditory cortex. Moreover, because auditory reafference (i.e., sounds related to object lifting, replacement, etc.) is masked in our experiments both by the loud scanner noise and the headphones participant’s wear to protect their hearing (and allow the delivery of auditory commands), we are able to test the extent to which modulations in auditory cortex are automatic (i.e., are not contingent on whether auditory feedback is available or masked) and motor-related in origin (Schneider et al. 2014).

In the first of these experiments, in each trial we had participants first prepare, and then execute (after a jittered delay interval) either a left or right hand object lift-and-replace action, which were cued by two nonsense auditory commands (“Compty” or “Midwig”; see Fig 1). Importantly, halfway throughout each experimental run, participants were required to switch the auditory command-to-hand mapping (i.e., if “Compty” cued a left hand object lift-and-replace action in the first half of the experimental run, then “Compty” would cue a right hand object lift-and-replace action in the second half of the experimental run; see Fig 1B). Critically, this design allowed us to examine early auditory cortex activity during the planning of two distinct hand actions (left versus right hand movements), with invariance to the actual auditory commands (e.g., “Compty”) used to instruct those hand actions. In the second of these experiments, in each trial we had participants first prepare, and then execute (after a fixed delay interval) either a right hand object lift-and-replace action or a target-directed eye movement (see Fig. 4). Unlike in Experiment 1, both of these movements were instructed via change in the colour of the central fixation light, thus allowing us to examine early auditory cortex activity during the planning of two distinct effector movements (hand versus eye) in the absence of any direct auditory input (i.e., no auditory commands). As such, any neural differences in the auditory cortex prior to movement in both of these experiments is likely to reflect modulations related to motor, and not bottom-up sensory, processing.

**Figure 1.**
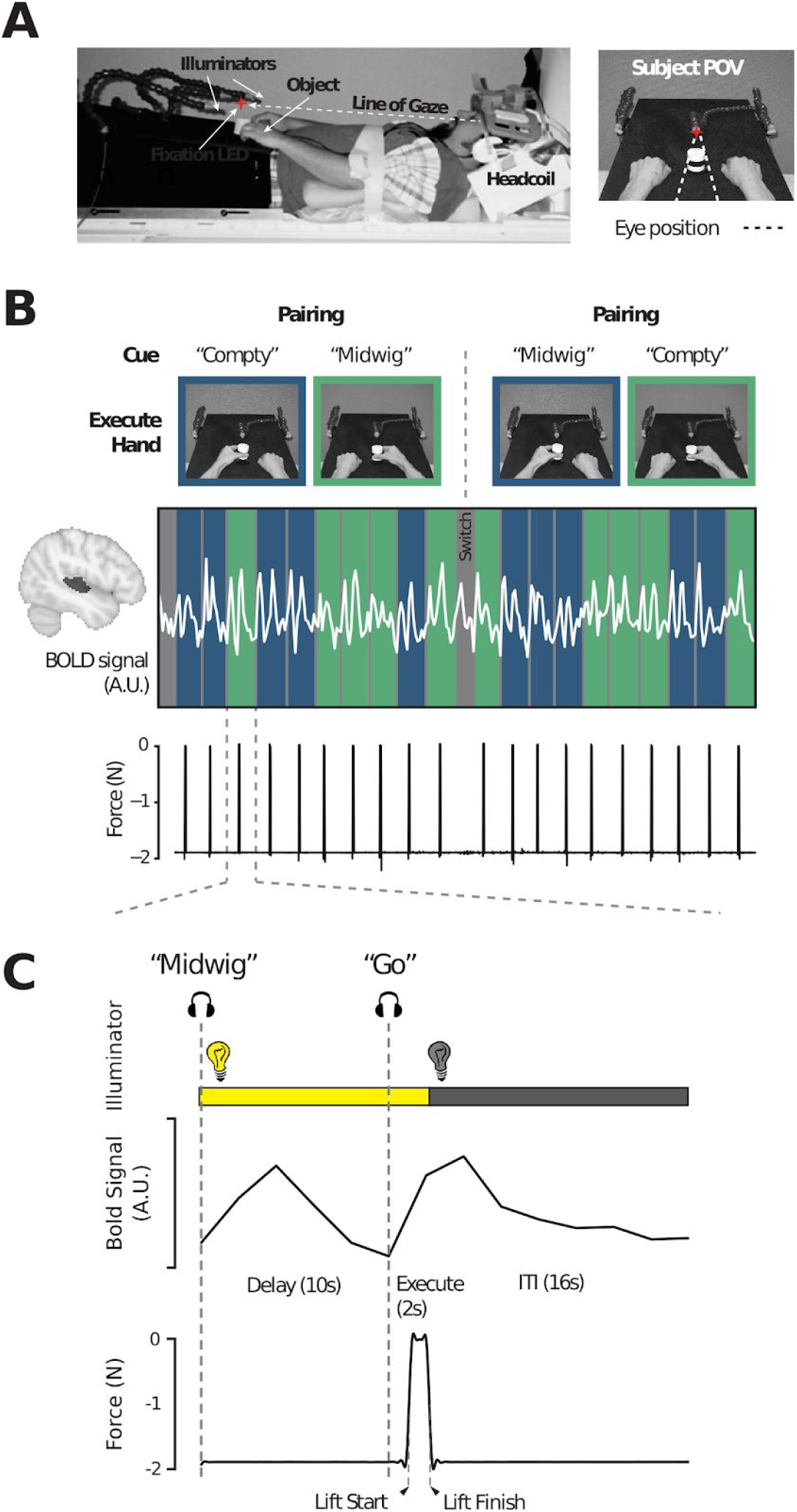
Experiment 1 setup and task overview. **A.** MRI setup (left) and subject point-of-view (right) of the experimental workspace. Red star indicates the fixation LED placed above the object. Illuminator LEDs, attached to the flexible stalks, are shown on the left and right. **B.** Example fMRI run of 20 task trials. Color-coded columns (blue = left hand, green = right hand) demarks each trial and the associated time-locked BOLD activity from superior temporal gyrus (STG; shaded in dark gray on cortex, left) of an exemplar subject is indicated by the overlaid white trace. Pairings between auditory cue (“Compty” or “Midwig”) and hand (left or right) are indicated in the pictures above, and were reversed halfway through each run following a ‘Switch’ auditory cue, such that each hand is paired with each auditory cue in every experimental run (see Methods). The corresponding force sensor data, used to track object lifting, is shown below. **C.** Sequence of events and corresponding single-trial BOLD and force sensor data of an exemplar trial from a representative participant in which ‘Midwig’ cued a right-handed movement. Each trial begins with the hand workspace being illuminated while, simultaneously, participants receive the auditory cue (“Compty” or “Midwig”). This is then followed by a jittered 6-12s Delay interval (10s in this exemplar trial). Next, an auditory “Go” cue initiates the start of the 2s Execute epoch, during which the subject grasp-and-lifts the object (shown by the force trace; arrows indicate the start of the lift and object replacement). Following the 2 s Execute epoch, illumination of the workspace is extinguished and subjects then wait a fixed 16s intertrial interval (ITI) prior to onset of the next trial. See also Supplemental Fig. 1 for a more detailed overview of the trial sequence and the data obtained from a separate behavioural training session.

### Participants

Sixteen healthy right-handed subjects (8 females, 21-25 years of age) participated in Experiment 1, which involved one behavioural testing session followed by two fMRI testing sessions (a localizer testing session, and then the experimental testing session, performed on separate days approximately 1-2 weeks apart). A separate cohort of fifteen healthy right-handed subjects (8 females, 20-32 years of age) participated in Experiment 2 approximately a year after Experiment 1, which involved one behavioural testing session and one fMRI testing session. Right-handedness was assessed with the Edinburgh handedness questionnaire (Oldfield 1971). Informed consent and consent to publish were obtained in accordance with ethical standards set out by the Declaration of Helsinki (1964) and with procedures cleared by the Queen’s University Health Sciences Research Ethics Board. Subjects were naïve with respect to the hypotheses under evaluation and received monetary compensation for their involvement in the study. Data from one subject in Experiment 1 and from two subjects in Experiment 2 were excluded from further analyses due to data collection problems in the experimental testing sessions, resulting in final sample sizes of 15 and 13 subjects, respectively. Meanwhile, all 16 subjects from Experiment 1 were used for the localizer testing session.

### Experiment 1

#### Experimental apparatus

The experimental setup for both the localizer and experimental testing sessions was a modification from some of our previous fMRI studies (Gallivan et al. 2011; Gallivan, Chapman, et al. 2013; Hutchison and Gallivan 2018), and consisted of a black platform placed over the waist and tilted away from the horizontal at an angle (~15°) to maximize comfort and target visibility. The MRI head coil was tilted slightly (~20°) and foam cushions were used to give an approximate overall head tilt of 30°. To minimize limb-related artifacts, subjects had the right and left upper-arms braced, limiting movement of the arms to the elbow and thus creating an arc of reachability for each hand. The exact placement of object stimuli on the platform was adjusted to match each subject’s arm length such that all required actions were comfortable and ensured that only movement of the forearm, wrist and fingers was required. The platform was illuminated by two bright white Light Emitting Diodes (LEDs) attached to flexible plastic stalks (Loc-Line, Lockwood Products, Lake Oswego, OR, USA) located to the left and right of the platform. To control for eye movements, a small red fixation LED, attached to a flexible plastic stalk, was positioned above the hand workspace and located ~5 cm beyond the target object position (such that the object appeared in the subject’s lower visual field). Experimental timing and lighting were controlled with in-house software created with C++ and MATLAB (The Mathworks, Natnick, MA). Throughout fMRI testing, the subject’s hand movements were monitored using an MR-compatible infrared-sensitive camera (MRC Systems GmbH, Heidelberg, Germany), optimally positioned on one side of the platform and facing towards the subject. The videos captured during the experiment were analyzed offline to verify that the subjects were performing the task as instructed and to identify error trials.

#### Auditory Localizer Task

A separate, block-design localizer task was collected to independently identify auditory cortex and higher-order language regions of interest (ROIs) for use in the analyses of main experimental task. This auditory localizer task included three conditions: (1) Intact speech trials (Intact), which played one of 8 unique excerpts of different speeches; (2) scrambled speech trials (Scrambled), which were incoherent signal-correlated noise versions of the speech excerpts (i.e. applying an amplitude envelope of the speech to uniform Gaussian white noise, ensuring that the noise level was utterance-specific and exactly intense enough at every moment to mask the energy of the spoken words); and (3) rest trials (Rest), in which no audio was played (subjects thus only heard background MRI scanner noise during this time). Trials lasted 20 s each and alternated, in pseudo-random order, between Intact Speech, Scrambled Speech, and Rest for a total of 24 trials in each experimental run. In addition, a 20 s baseline block was placed at the beginning of each experimental run. Each localizer run totaled 500 s and participants completed 2 of these runs during testing (resulting in 16 repetitions per experimental condition per subject). To encourage that participants maintained attention throughout this auditory localizer run, they were required to monitor each of the Intact speeches and let the experimenter know, following the run, whether any of them were repeated. This repeat happened in only one of the experimental runs and each and every subject correctly identified the repeat and non-repeat run (100% accuracy).

#### Motor Localizer task

Four experimental runs of a motor localizer task were also collected alongside the auditory localizer task, which constituted a block-design protocol that alternated between subtasks designed to localize eight separate motor functions. Task set up and details for all eight conditions are described in Supplemental Fig. 2. The hand grasping condition from this localizer task was used to define dorsal premotor cortex (PMd), which we used as a basis for comparison with our auditory cortex decoding findings (see Results).

The motor and auditory localizer testing session lasted approximately 2 hours and included set-up time, one 7.5 minute high-resolution anatomical scan and 6 functional scanning runs, wherein subjects alternated between performing two runs of the motor localizer task and one run of the auditory localizer, twice. A brief (~10 minute) practice session was carried out before the localizer testing session in the MRI control room in order to familiarize participants with localizer tasks.

#### Main Experimental Task

In our experimental task (see Fig. 1), we used a delayed movement paradigm wherein, on each individual trial, subjects were first auditorily cued (via headphones) to prepare either a left vs. right hand object grasp-and-lift action on a centrally located cylindrical object (1.9 N weight). Then, following a variable delay period, they were prompted to execute the prepared hand action (Gallivan, McLean, Flanagan, et al. 2013; see also Gallivan, McLean, Valyear, et al. 2013). At the start of each event-related trial (see Fig. 1C), simultaneously with the LED illuminator lights going on (and the subject’s workspace being illuminated), subjects received one of two nonsense speech cues, “Compty” or “Midwig”. For a given trial, each nonsense speech cue was paired with a corresponding hand action (e.g., subjects were instructed that, for a predetermined set of trials, “Compty” cued a left hand movement whereas “Midwig” cued a right hand movement). [*Note that nonsense speech commands were chosen because semantically meaningful words such as “left” and “right” would already have strong cognitive associations for participants*.] Following the delivery of the auditory command, there was a jittered delay interval of 6-12 s (a Gaussian random jitter with a mean of 9 s), after which the verbal auditory command “Go” was delivered, prompting subjects to execute the prepared grasp-and-lift action. For the execution phase of the trial, subjects were required to precision grasp-and-then-lift the object with their thumb and index finger (~2 cm above the platform, via a rotation of the wrist), hold it in midair for ~1 s, and then replace it. Subjects were instructed to keep the timing of each hand action as similar as possible throughout the study. Two seconds following the onset of this “Go” cue, the illuminator lights were extinguished, and subjects then waited 16 s for the next trial to begin (intertrial interval, ITI). Throughout the entire time course of the trial, subjects were required to maintain gaze on the fixation LED.

These event-related trials were completed in two separate blocks per experimental run. At the beginning of each experimental run, the experimenter informed subjects of the auditory-hand mapping to be used for the first 10 event-related trials of the experimental run (e.g. “Compty” for left hand (left hand) movements, “Midwig” for right hand (right hand) movements; 5 intermixed trials of each type). After the 10th trial, the illuminator was turned on (for a duration of 6 s) and subjects simultaneously heard the auditory command “Switch” (following by a 16 s delay). This indicated that the auditory-hand mapping would now be reversed for the remaining 10 event-related trials (i.e., “Compty” would now cue a right hand movement whereas “Midwig” would now cue a left hand movement). The sequential ordering of this auditory-hand mapping was counterbalanced across runs, and resulted in a total of 4 different auditory-hand mappings (and thus, trial types) per experimental run: Compty-left hand, Compty-right hand, Midwig-left hand, and Midwig-right hand (with 5 repetitions each; 20 trials in total per run). With the exception of the blocked nature of these trials, these trial types were pseudorandomized within a run and counterbalanced across all runs so that each trial type was preceded and followed equally often by every other trial type across the entire experiment.

Separate practice sessions were carried out before the actual fMRI experiment to familiarize subjects with the delayed timing of the task. One of these sessions was conducted before subjects entered the scanner (See *Behavioural Control Experiment* below) and another was conducted during the anatomical scan (collected at the beginning of the experimental testing session). The experimental testing session for each subject lasted approximately 2 hours and included set-up time, one 7.5 minute high-resolution anatomical scan (during which subjects could practice the task) and eight functional scanning runs (for a total of 160 trials; 40 trials for each auditory-motor mapping). Each functional run (an example run shown in Fig. 1B) had a duration of 576 s, with a 30-60 s break in between each run. Lastly, a resting state functional scan, in which subjects lay still (with no task) and only maintained gaze on the fixation LED, was performed for 12 minutes (*data not analyzed here*).

During MRI testing, we also tracked subjects’ behaviour using an MRI-compatible force sensor located beneath the object (Nano 17 F/T sensors; ATI Industrial Automation, Garner, NC), and attached to our MRI platform. This force sensor, which was capped with a flat circular disk (diameter of 7.5 cm), supported the object. The force sensor measured the vertical forces exerted by the object (signals sampled at 500 Hz and low-pass filtered using a 5^th^ order, zero-phase lag Butterworth filter with a cutoff frequency of 5 Hz), allowing us to track both subject reaction time (RT), which we define as the time from the onset of the “Go” cue to object contact (Mean = 1601ms, SD = 389ms), and movement time (MT), which we define as the time from object lift to replacement (Mean = 2582ms, SD = 662ms), as well as generally monitor task performance. Note that we did not conduct eye tracking during this or any of the other MRI scan sessions because of the difficulties in monitoring gaze in the head-tilted configuration with standard MRI-compatible eye trackers (due to occlusion from the eyelids)(Gallivan et al. 2014, 2016, 2019a).

### Experiment 2

This study and experimental setup was similar to Experiment 1, with the exception that: (1) participants performed either a right hand object grasp-and-lift action on a centrally located cylindrical object (1.9 N weight) or a target-directed eye movement towards that same object (i.e., two experimental conditions), (2) the Delay epoch was a fixed duration (12 s), and (3) subjects were cued about the upcoming movement to be executed via a 0.5 s change in the fixation LED colour (from red to either blue or green, with the colour-action mapping being counterbalanced across subjects; i.e., a LED change to blue cued a grasp action in half the subjects, and cued an eye movement in the other half of subjects). The eye movement action involved the subject making a saccadic eye movement from the fixation LED to the target object, holding that position until the illuminator LEDs were extinguished, and then returning their gaze to the fixation LED. The two trial types, with 5 repetitions per condition (10 trials total), were pseudorandomized as in Experiment 1. Each subject participated in at least eight functional runs.

### Data Acquisition and Analysis

Subjects were scanned using a 3-Tesla Siemens TIM MAGNETOM Trio MRI scanner located at the Centre for Neuroscience Studies, Queen’s University (Kingston, Ontario, Canada). An identical imaging protocol was used for both Experiments 1 and 2, with the exception of slice thickness (Experiment 1 = 4mm; Experiment 2 = 3mm). In both experiments, MRI volumes were acquired using a T2*-weighted single-shot gradient-echo echo-planar imaging acquisition sequence (time to repetition = 2000 ms, in-plane resolution = 3 mm x 3 mm, time to echo = 30 ms, field of view = 240 mm x 240 mm, matrix size = 80 x 80, flip angle = 90°, and acceleration factor (integrated parallel acquisition technologies, iPAT) = 2 with generalized auto-calibrating partially parallel acquisitions reconstruction). Each volume comprised 35 contiguous (no gap) oblique slices acquired at a ~30° caudal tilt with respect to the plane of the anterior and posterior commissure (AC-PC). Subjects were scanned in a head-tilted configuration, allowing direct viewing of the hand workspace. We used a combination of imaging coils to achieve a good signal to noise ratio and to enable direct object workspace viewing without mirrors or occlusion. Specifically, we tilted (~20° degrees) the posterior half of the 12-channel receive-only head coil (6-channels) and suspended a 4-channel receive-only flex coil over the anterior-superior part of the head. An identical T1-weighted ADNI MPRAGE anatomical scan was also collected for both Experiments 1 and 2 (time to repetition = 1760 ms, time to echo = 2.98 ms, field of view = 192 mm x 240 mm x 256 mm, matrix size = 192 x 240 x 256, flip angle = 9°, 1 mm isotropic voxels).

### fMRI data preprocessing

Preprocessing of functional data collected in the localizer testing session, and Experiments 1 and 2, was performed using *fMRIPrep* 1.4.1 (Esteban et al. 2018), which is based on *Nipype* 1.2.0 (Gorgolewski et al. 2011; Esteban et al. 2019).

#### Anatomical data preprocessing

The T1-weighted (T1w) image was corrected for intensity non-uniformity (INU) with N4BiasFieldCorrection (Tustison et al. 2010), distributed with ANTs 2.2.0 (Avants et al. 2008), and used as T1w-reference throughout the workflow. The T1w-reference was then skull-stripped with a Nipype implementation of the antsBrainExtraction.sh workflow (from ANTs), using OASIS30ANTs as target template. Brain tissue segmentation of cerebrospinal fluid (CSF), white-matter (WM) and gray-matter (GM) was performed on the brain-extracted T1w using fast (FSL 5.0.9, (Zhang et al. 2001)). Brain surfaces were reconstructed using recon-all (FreeSurfer 6.0.1, (Dale et al. 1999)), and the brain mask estimated previously was refined with a custom variation of the method to reconcile ANTs-derived and FreeSurfer-derived segmentations of the cortical gray-matter of Mindboggle (Klein et al. 2017). Volume-based spatial normalization to standard space (voxel size = 2 × 2 × 2 mm) was performed through nonlinear registration with antsRegistration (ANTs 2.2.0), using brain-extracted versions of both T1w reference and the T1w template. The following template was selected for spatial normalization: FSL’s MNI ICBM 152 non-linear 6th Generation Asymmetric Average Brain Stereotaxic Registration Model [(Evans et al. 2012); TemplateFlow ID: MNI152NLin6Asym].

#### Functional data preprocessing

For each BOLD run per subject (across all tasks and/or sessions), the following preprocessing was performed. First, a reference volume and its skull-stripped version were generated using a custom methodology of fMRIPrep. The BOLD reference was then co-registered to the T1w reference using bbregister (FreeSurfer) which implements boundary-based registration (Greve and Fischl 2009). Co-registration was configured with nine degrees of freedom to account for distortions remaining in the BOLD reference. Head-motion parameters with respect to the BOLD reference (transformation matrices, and six corresponding rotation and translation parameters) are estimated before any spatiotemporal filtering using mcflirt (FSL 5.0.9, (Jenkinson et al. 2002)). BOLD runs were slice-time corrected using 3dTshift from AFNI 20160207 (Cox and Hyde 1997). The BOLD time-series were normalized by resampling into standard space. All resamplings were performed with a single interpolation step by composing all the pertinent transformations (i.e. head-motion transform matrices, and co-registrations to anatomical and output spaces). Gridded (volumetric) resamplings were performed using antsApplyTransforms (ANTs), configured with Lanczos interpolation to minimize the smoothing effects of other kernels (Lanczos 1964).

Many internal operations of fMRIPrep use Nilearn 0.5.2 (Abraham et al. 2014), mostly within the functional processing workflow. For more details of the pipeline, see the section corresponding to workflows in fMRIPrep’s documentation.

#### Post-processing

Additional post-processing was performed for specific analyses. Normalized functional scans were temporally filtered using a high-pass filter (cutoff = 0.01 Hz) to remove low-frequency noise (e.g. linear scanner drift), either as part of GLMs (see below) or directly (as in time-point decoding analyses). For the localizer data, normalized functional scans were spatially smoothed (6mm FWHM Gaussian kernel; SPM12) prior to GLM estimation to facilitate subject overlap. [Note that no spatial smoothing was performed on the experimental task data sets, wherein multi-voxel pattern analyses were performed.]

#### Error trials

Error trials were identified offline from the videos recorded during the experimental testing session and were excluded from analysis by assigning these trials predictors of no interest. Error trials included those in which the subject performed the incorrect instruction (Experiment 1: 9 trials, 4 subjects; Experiment 2: 1 trial, 1 subject) or contaminated the delay phase data by slightly moving their limb or moving too early (Experiment 1: 7 trials, 4 subjects; Experiment 2: 1 trial, 1 subject). The fact that subjects made so few errors when considering the potentially challenging nature of the tasks (e.g., in Experiment 1 having to remember whether “Compty” cued a left hand or right hand movement on the current trial) speaks to the fact that subjects were fully engaged during experimental testing and very well practiced at the task prior to participating in the experiment.

### Statistical Analyses

#### General Linear Models

For the localizer task analyses, we carried out subject-level analysis using SPM12’s first-level analysis toolbox to create general linear models (GLM) for each task (auditory and motor). Each GLM featured condition predictors created from boxcar functions convolved with a double-gamma hemodynamic response function (HRF), which were aligned to the onset of each action/stimulus block with durations dependent on block length (i.e., 10 imaging volumes for both localizer tasks). Temporal derivatives of each predictor and subjects’ six motion parameters obtained from motion correction were included as additional regressors. The Baseline/Fixation epochs were excluded from the model; therefore all regression coefficients (betas) were defined relative to the baseline activity during these time points.

In the experimental tasks, we employed a Least-Squares Separate procedure (Mumford et al., 2012) to extract beta coefficient estimates for decoding analyses. This procedure generated separate GLM models for each individual trial’s Delay and Execute epochs (e.g., In Experiment 1: 20 trials x 2 epochs x 8 runs = 320 GLMs). The regressor of interest in each model consisted of a boxcar regressor aligned to the start of the epoch of interest. The duration of the regressor was set to the duration of the cue that initiates the epoch (0.5s): the auditory command cue (‘Compty’ or ‘Midwig’) and the visual cue (fixation LED colour change) for the Delay epoch in Experiment 1 and 2, respectively; and the auditory ‘Go’ cue for the Execute epoch in both experiments. For each GLM, we included a second regressor that comprised of all remaining trial epochs in the experimental run. Each regressor was then convolved with a double-gamma HRF, and temporal derivatives of both regressors were included along with subjects’ six motion parameters obtained from motion correction. Isolating the regressor of interest in this single-trial fashion reduces regressor collinearity, and has been shown to be advantageous in estimating single-trial voxel patterns and for multi-voxel pattern classification (Mumford et al. 2012).

#### Region of interest (ROI) selection

Regions of interests (ROI) were identified based on second-level (group) analyses of first-level contrast images from each subject. Early auditory cortex ROIs were identified for both Experiment 1 and 2 by thresholding a Scrambled Speech > Rest group contrast at an uncorrected voxelwise threshold of p < 10^-5^. This procedure identified tight superior temporal gyrus (STG) activation clusters in left and right Heschl’s gyrus (HG), the anatomical landmark for primary (core) auditory cortex (Morosan et al. 2000, 2001; Rademacher et al. 2001; Da Costa et al. 2011), and more posteriorly on the superior temporal plane (Planum Temporale, PT). We verified these locations by intersecting region masks for HG and PT obtained from the Harvard-Oxford anatomical atlas (Desikan et al. 2006) with the masks of left and right STG clusters. This allowed us to define, for each participant, voxels that were active for sound that fell in anatomically defined HG and PT. We considered HG and PT separately since they are at different stages of auditory processing: HG is the location of the core, whereas the PT consists of belt and probably parabelt regions, as well as possibly other types of cortical tissue (Hackett et al. 2014). Since our PT activity is just posterior to HG, we suspect that this is probably in belt or parabelt cortex, one or two stages of processing removed from core. Lastly, a more expansive auditory and speech processing network was obtained using a Intact Speech > Rest contrast with an uncorrected height threshold of *p* < .001 and cluster-extent correction threshold of p < .05. Together, these were used as three-dimensional binary masks to constrain our analyses and interpretations of motor-related effects in the auditory system.

#### Multi-voxel Pattern Analysis (MVPA)

For the experimental task, MVPA was performed with in-house software using Python 3.7.1 with Nilearn v0.6.0 and Scikit-Learn v0.20.1 (Abraham et al. 2014). All analyses implement support vector machine (SVM) binary classifiers (libSVM) using a radial-basis function kernel and with fixed regulation parameter (C = 1) and gamma parameter (automatically set to the reciprocal of the number of voxels) in order to compute a hyperplane that best separated the trial responses. The pattern of voxel beta coefficients from the single-trial GLMs, which provided voxel patterns for each trial’s Delay and Execute epochs, were used as inputs into the binary classifiers. These values were standardized across voxels such that each voxel pattern had a mean of 0 and standard deviation of 1. Therefore, the mean univariate signal for each pattern was removed in the ROI.

Decoding accuracies for each subject were computed as the average classification accuracy across train-and-test iterations using a ‘leave-one-run-out’ cross-validation procedure. This procedure was performed separately for each ROI, trial epoch (Delay and Execute), and pairwise discrimination (left hand vs right hand movements and “Compty” vs “Midwig” in Experiment 1; and hand vs. eye movements in Experiment 2). We statistically assessed decoding significance at the group-level using one-tailed t-tests vs. 50% chance decoding. To control for the problem of multiple comparisons within each ROI (i.e. the number of pairwise comparisons tested), we applied a Benjamini-Hochberg false-discovery rate (FDR) correction of q<0.05.

Significant decoding accuracies were further verified against null distributions constructed by a two-step permutation approach based on (Stelzer et al. 2013). In the first step, distributions of chance decoding accuracies were independently generated for each subject using 100 iterations. In each iteration, class labels were randomly permuted within each run, and a decoding accuracy was computed by taking the average classification accuracy following leave-one-run-out cross validation. In the second step, the subject-specific distributions of decoding accuracies were used to compute a distribution of 10,000 group-average decoding accuracies. Here, in each iteration, a decoding accuracy was randomly selected from each subject distribution and the mean decoding accuracy across subjects was calculated. The distribution of group-average decoding accuracies was then used to compute the probability of the actual decoding accuracy.

### Searchlight Pattern-Information Analyses

To complement our MVPA ROI analyses in both Experiments 1 and 2, we also performed a pattern analysis in each subject using the searchlight approach (Kriegeskorte et al. 2006). Given the scope of this paper (i.e., to examine the top-down modulation of auditory cortex during planning), we constrained this searchlight analysis to the auditory network mask defined by the Intact Speech > Rest contrast using the independent auditory localizer data (allowing us to localize both lower-order and higher-order auditory regions). In this procedure, the SVM classifier moved through each subjects’ localizer-defined auditory network in a voxel-by-voxel fashion whereby, at each voxel, a sphere of surrounding voxels (radius of 4mm; 33 voxels) were extracted, z-scored within pattern (see above), and input into the SVM classifier. The decoding accuracy for that sphere of voxels was then written to the central voxel. This searchlight procedure was performed separately with beta coefficient maps for the Delay and Execute epochs based on the GLM procedure described above, which yielded separate Delay and Execute whole-brain decoding maps. To allow for group-level analyses, the decoding maps were smoothed (6mm FWHM Gaussian kernel) in each subject, which is particularly pertinent given the known neuroanatomical variability in human auditory cortex (Hackett et al. 2001; Zoellner et al. 2019; Ren et al. 2020). Then, for each voxel, we assessed statistical significance using a one-tailed t-test versus 50% chance decoding.

Group-level decoding maps for Delay and Execute epochs were thresholded at p < .01 and cluster corrected to p < .05 using Monte-Carlo style permutation tests with AFNI’s 3dClustSim algorithm (Cox 1996; Cox et al. 2017). Cluster correction was additionally validated using a nonparametric approach (see Supplemental Fig. 3). We note that this latter approach resulted in more liberal cluster correction thresholds in comparison with 3dClustSim; we therefore opted to use the more conservative thresholds obtained from 3dClustSim.

## RESULTS

### Experiment 1

#### Delay period decoding of hand information from early auditory cortex

To determine whether signals related to hand movement planning influence early auditory cortex activity, we extracted the trial-related voxel patterns (beta coefficients) associated with the Delay (as well as Execute) epochs from early auditory cortex. To this end, we first functionally identified, using data from an independent auditory localizer task (see Methods), fMRI activity in the left and right superior temporal gyrus (STG). To provide greater specificity with regards to the localization of potential motor planning-related effects, we further delineated these STG clusters based on their intersections with Heschl’s gyrus (HG) and the Planum Temporale (PT), two adjacent human brain regions associated with primary and higher-order cortical auditory processing, respectively (Poeppel et al. 2012)(see Fig. 2A,B for our basic approach). Next, for each of these 3 regions (STG, and its subdivisions into HG and PT) we used their z-scored Delay epoch voxel activity patterns as inputs to a support vector machine (SVM) binary classifier. In order to derive main-effects of hand information (i.e., examine decoding of upcoming left hand vs. right hand movements) versus auditory cue information (examine decoding of “Compty” vs. “Midwig” cues) and to increase the data used for classifier training, we performed separate analyses wherein we collapsed across auditory cue or hand trials, respectively. Our logic is that, when collapsing across auditory cue (i.e., re-labelling all trials based on the hand used), if we can observe decoding of hand information in auditory cortex during the Delay phase (prior to movement), then this information is represented with invariance to the cue, and thus sensory input (and vice versa).

**Figure 2.**
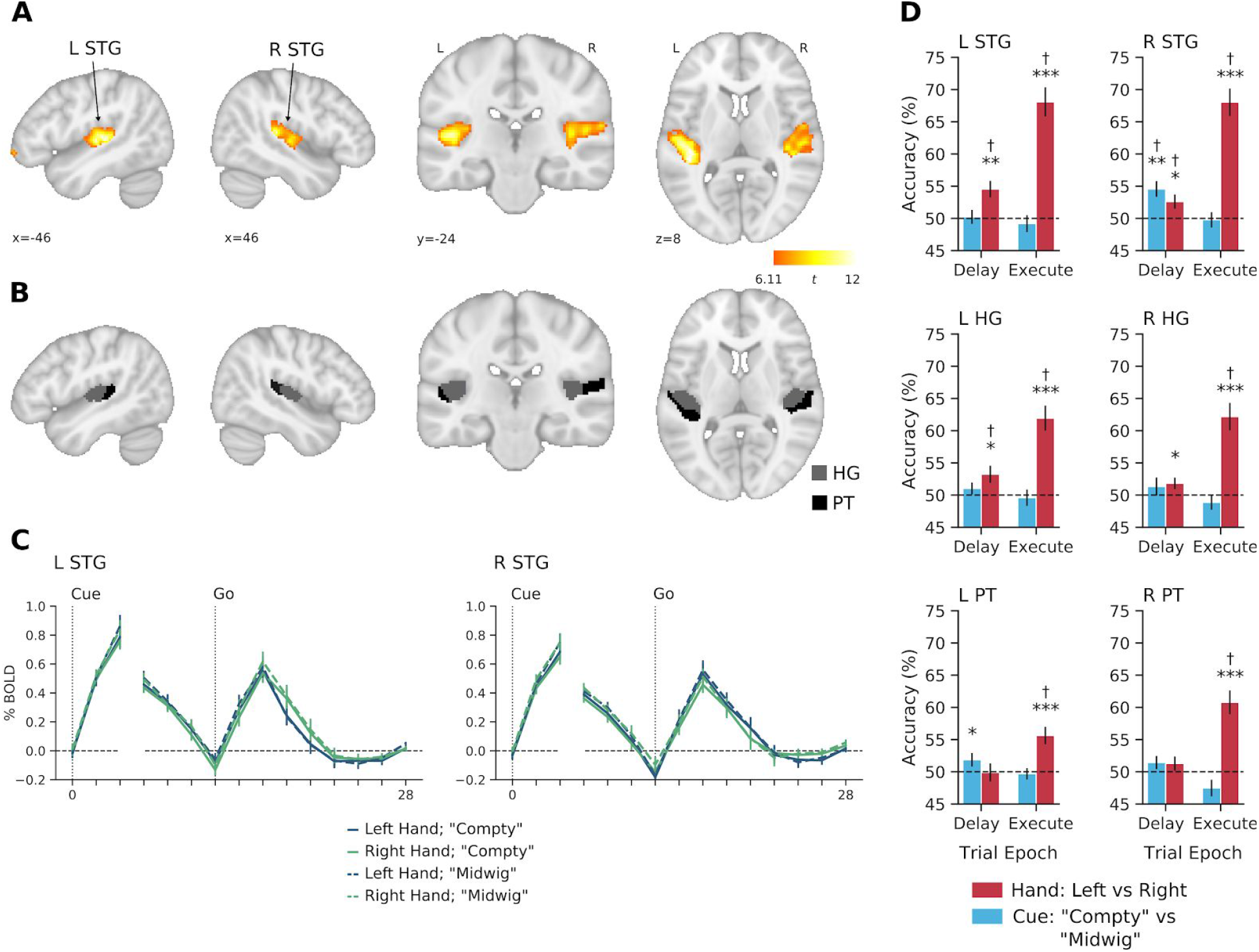
Decoding of hand information in early auditory cortex for Experiment 1. **A.** Left and right STG clusters defined at the group-level (N = 16) with the independent auditory localizer task using the contrast of Scrambled Speech > Rest (see Methods). **B.** Delineation of Heschl’s gyrus (HG) and Planum Temporale (PT), within each STG cluster from A. **C.** Group-averaged percent-signal change BOLD time courses for each trial type in left and right STG. The first three timepoints are time-locked to the onset of the Delay epoch and remaining time points are time-locked to onset of the Execute epoch. There is a high degree of overlap amongst the time courses for the different trial types. Note the separate increases in activity associated with delivery of the auditory cue (“Compy” or “Midwig”) and “Go” signal. **D.** Decoding accuracies for hand (red) and cue (blue) information. Hand and cue decoding accuracies were analyzed separately in each epoch (Delay and Execute) using one-sample t-tests (one-tailed) against chance level (50%). Error bars show ± 1 SE of mean. * p < .05, ** p < .01, *** p < .001, † FDR corrected q < .05.

Our analysis on the resulting classification accuracies (see Fig. 2D) revealed that information related to the upcoming hand actions to be performed (i.e., during the Delay epoch) was present in bilateral STG (left: t_14_ = 3.55, p = .002; right: t_14_ = 2.34, p = .017) and left HG (t_14_ = 2.43, p = .014). A significant effect was also found in right HG but it did not survive FDR correction (t_14_ = 2.06, p = .029). These findings were additionally confirmed using follow-up permutation analyses (see Supplemental Fig. 4A). Meanwhile, no significant decoding was found in left (t_14_ = −.074, p = .529) or right (t_14_ = 1.17, p = .131) PT. By contrast, during the Execute epoch, we found that hand decoding was robust in all three areas in both hemispheres (all p < .001, see Fig. 2D; all p < .001 in permutation analyses). Because our task did not pair the execution of hand movements to sound generation, and subjects would not have heard the auditory consequences associated with movement (e.g., object lifting and replacement) due to their wearing of headphones and the loud background noise of the scanner, these Execution results suggest that the modulation of auditory cortex activity is automatic and motor-related in nature (Schneider et al. 2014). Importantly, our finding of decoding in bilateral STG during the Delay period further suggests that this automatic and motor-related modulation also occurs well before execution, during movement preparation. An additional behavioural control experiment, performed prior to MRI testing (see Supplementary Material), suggests that the emergence of these hand-related effects are unlikely to be driven by systematic differences in eye position across trials (Werner-Reiss et al. 2003), since our trained participants exhibited highly stable fixations throughout the task (Supplemental Fig. 1).

In contrast to our motor-related hand decoding results, our analysis on the resulting classification accuracies for the sensory-related auditory cue (“Compty” vs. “Midwig”) revealed that, during the Delay epoch, information related to the delivered verbal cue was present only in right STG (t_14_ = 3.71, p = .001, see Fig. 2D). Left PT also showed significant decoding (t_14_ = 1.79, p = .048), although this did not survive FDR correction. Subsequent permutation analyses replicated this pattern of results (see Supplemental Fig. 4B). No cue decoding was found in the remaining ROIs (all p > .10). Critically, consistent with the fact that this auditory cue information was delivered to participants only during the Delay epoch (i.e., participants *always* received a “Go” cue at the Execute epoch, regardless of trial identity), we also observed no evidence of cue decoding during the Execute epoch (all p > .10).

#### Delay period decoding from auditory cortex mirrors that found in the motor system

To provide a basis for comparing and interpreting the above hand-related decoding effects in the auditory cortex, we also used the data from our experimental task to examine Delay epoch decoding in a positive control region, the dorsal premotor cortex (PMd). This region, which we independently identified PMd using a separate motor localizer task in our participants (see Methods), is well known to differentiate limb-related information during movement planning in both humans and nonhuman primates (Cisek et al. 2003; Gallivan, McLean, Flanagan, et al. 2013). As shown in Supplemental Fig. 2C, we found a remarkably similar profile of limb-specific decoding in this motor-related region to that observed in the auditory cortex (area STG). In summary, this PMd-result allows for two important observations. First, similar levels of action-related information can be decoded from early auditory cortex as from dorsal premotor cortex, the latter area known to have a well-established role in motor planning (Weinrich et al. 1984; Kaufman et al. 2010; Lara et al. 2018). Second, this Delay period decoding suggests that the representation of hand-related information evolves in a similar fashion prior to movement onset in both STG and PMd.

#### Searchlight analyses reveal the representation of hand information in early auditory cortex prior to movement

To complement our above ROI analyses, we also performed a group-level searchlight analysis within the wider auditory processing network, localized using our independent auditory localizer data (see Methods, Supplemental Fig. 5A). During the Delay epoch (see Fig. 3), two hand-related decoding clusters were identified in left STG, which includes a cluster centered on HG (212 voxels; peak, x=-48, y=-24, z=10, t_14_=8.31, p<.001) and a cluster spanning anterolateral portions of left STG (321 voxels; peak, x = −60, y = −2, z = −2, t_14_ = 5.72, p<.001). In the right hemisphere, one large cluster was revealed, which broadly spans across STG and superior temporal sulcus (519 voxels; peak, x = −50, y = −22, z = 8, t_14_ = 6.06, p<.001). Notably, when we examined cue-related decoding during the Delay epoch (i.e., decoding the auditory command ‘Compty’ vs. ‘Midwig’), we found one cluster in right STG (272 voxels; peak, x = 58, y = −6, z = 8, t_14_ = 8.27, p<.001), which did not overlap with the hand decoding clusters. This result suggests that, rather than a multiplexing of hand-related and cue-related within a common region of auditory cortex, separate subregions of the auditory system are modulated by motor-related (hand) versus sensory-related (auditory cue) information. In addition, the overlap of hand decoding clusters on bilateral HG and STG, as well as a cue decoding cluster in right STG, replicate our basic pattern of ROI-based results presented in Fig. 2.

**Figure 3.**
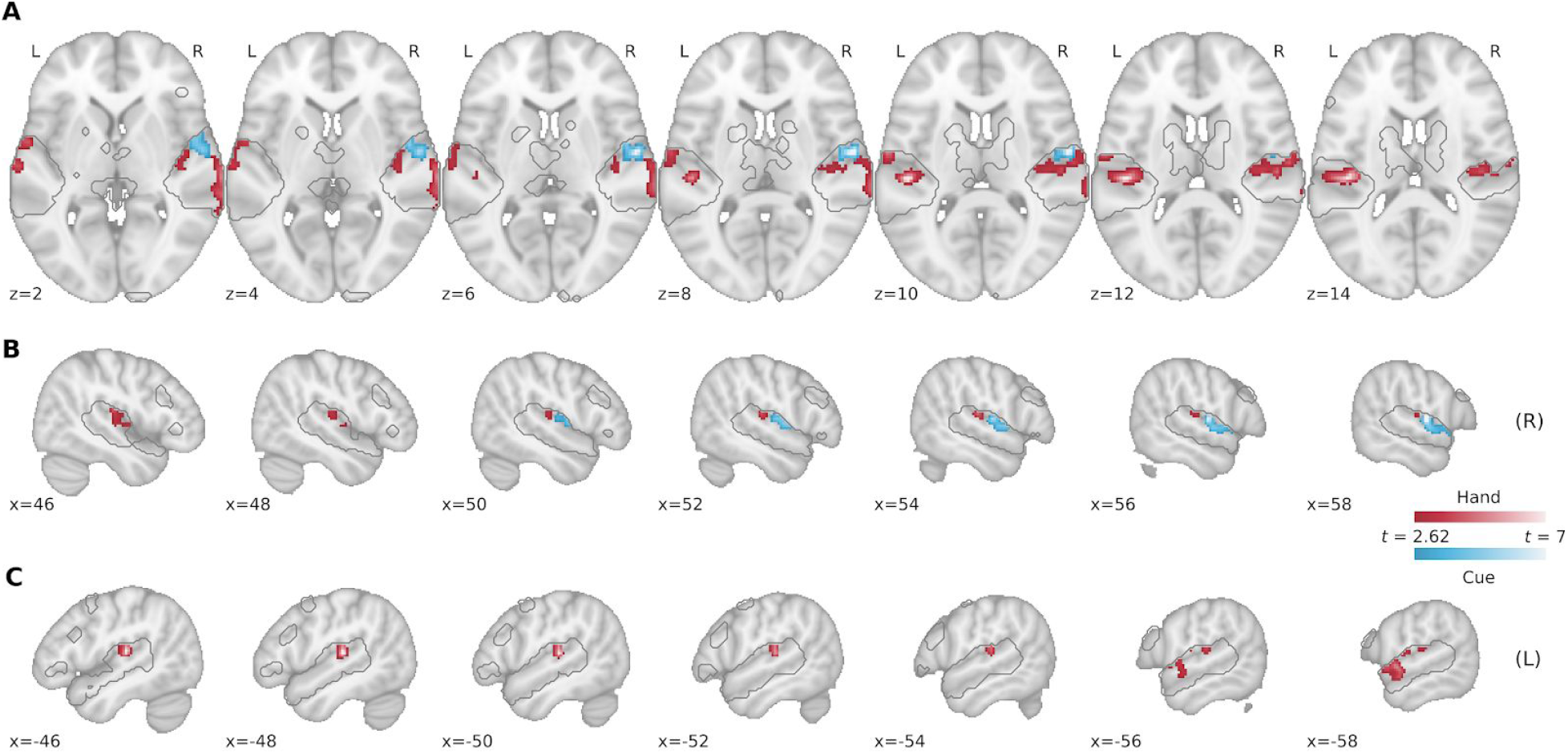
The representation of motor-related (hand) information during the Delay epoch in Experiment 1 is spatially distinct from the representation of sensory-related (auditory cue) information within superior temporal gyrus. Searchlight analyses were restricted to a mask defined by significant voxels in a Intact Speech > Rest contrast using an independent auditory localizer task (gray traced regions; see Methods). Group-level searchlight maps were thresholded at t > 2.62 (one-tailed p < .01) and cluster-corrected at p < .05. **A.** Transverse slices of significant decoding clusters for hand (red) and auditory cue (blue) in early auditory areas during the Delay epoch. **B.** Sagittal slices of the right hemisphere. **C.** Sagittal slices of left hemisphere.

A searchlight analysis using the Execute epoch data revealed a far more extensive pattern of hand decoding throughout the auditory network, with significant decoding extending all along the superior and middle temporal gyri bilaterally, and even into the basal ganglia and medial temporal cortex (see Supplemental Fig. 5B). By contrast, and in line with our ROI-based results, no cue decoding clusters were detected during the Execute epoch. These searchlight findings, when considered jointly with our ROI-based results, provide initial evidence that movement preparation selectively modulates neural activity patterns in early auditory cortex.

### Experiment 2

Our analyses from Experiment 1 demonstrate that hand-related information (left vs. right hand) is represented bilaterally in early auditory cortex well prior to movement. What remains undetermined from this first study, however, is the extent to which these modulations in auditory cortex purely reflect a motor-specific process. Indeed, one alternative explanation of the Experiment 1 results is that the decoding of limb-specific information could reflect some type of auditory working-memory and/or rehearsal process, whereby, during the delay period, participants translate the auditory command (e.g., ‘Compty’) into the corresponding auditory instruction (e.g., subvocalize to themselves “Right hand”). If true, then what we interpret here as ‘hand-related’ decoding may instead reflect a subvocalization process that recruits auditory cortex. Indeed, prior fMRI work has shown that the both the rehearsal (Paulesu et al. 1993) and maintenance (Kumar et al. 2016) of auditory information involves the activation of bilateral auditory cortex, and in Experiment 1, we are unable to disentangle such rehearsal/maintenance effects from the increase in BOLD activity related to the actual auditory cue delivered to participants at the onset of Delay epoch (“Compty” and “Midwig”; see Figure 2C). Thus, what is needed to disentangle these effects is a delayed movement task in which no auditory cues are used to instruct movement at the onset of the Delay epoch. This would allow us to examine whether we still observe an increase in BOLD activity during the delay period (as in Figure 2C), which would be consistent with the alternative explanation of our results that the ‘hand-related’ decoding observed instead reflects an auditory rehearsal/maintenance process.

To rule out the aforementioned potential confound and to replicate and extend the effects of Experiment 1 to different effector systems, in a second experiment we modified a classic task from primate neurophysiology used to dissociate motor-versus sensory-related coding during action planning (Snyder et al. 1997; Cui and Andersen 2007). In our version of this delayed movement task, in each trial, participants either grasped an object with their right hand or made an eye movement towards it (Fig. 4A). Importantly, unlike in Experiment 1 (wherein we used auditory commands to instruct actions at the onset of the Delay peoch), here we cued the two movements via a change in the colour of the fixation LED (Fig. 4A). As such, we could examine whether, in the absence of any auditory input at the onset of the Delay period, we still find increased activity in bilateral auditory cortex during the delay period. Such an observation would be consistent with an auditory rehearsal/maintenance process (Paulesu et al. 1993; Kumar et al. 2016). By contrast, if we were to instead find that the decoding of the upcoming action to be performed (i.e., hand vs. eye movement) occurs in the absence of any net change in auditory activity, then this would suggest that such decoding is linked to a top-down motor-related process.

**Figure 4.**
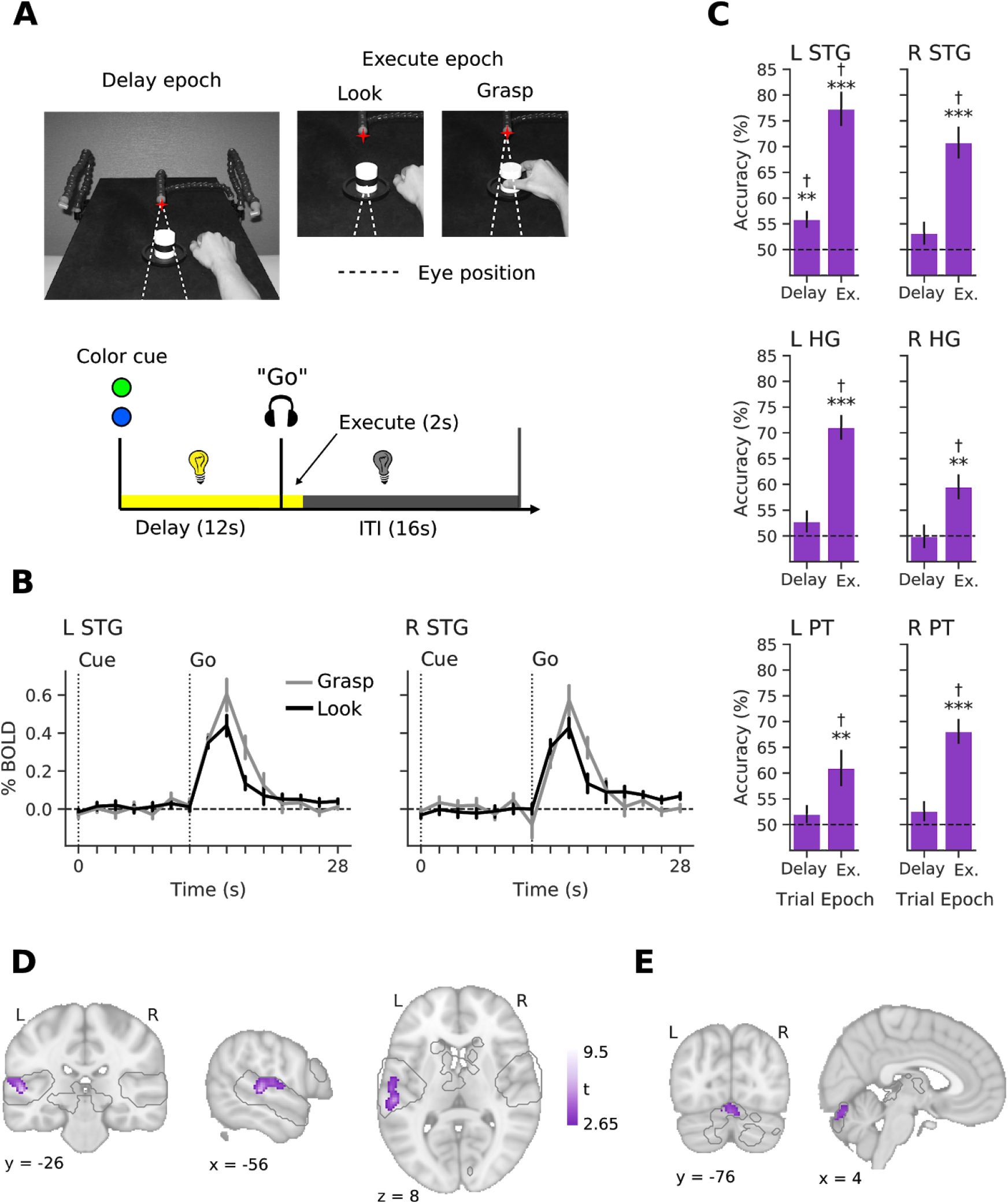
Experiment 2 shows effector-specific decoding in (contralateral) left auditory cortex and that it occurs despite baseline-levels of auditory activity during the delay period. **A.** Subject point of view of the experimental workspace in Delay and Execute epochs (top) and trial flow (bottom). Red star indicates the fixation illuminator LED. **B.** Group-averaged BOLD timecourses for the Grasp (grey) and Eye movement (‘Look’, black) conditions in left and right STG. Note the absence of any net change in BOLD activity during the delay epoch for both trial types. **C.** Decoding accuracies for Grasp vs Look conditions during the Delay and Execute epochs in all auditory ROIs. Decoding accuracies for each epoch were tested against chance decoding (50%) using one-sample t-tests. Error bars show ± 1 SE of mean. * p < .05, ** p < .01, *** p < .001, † FDR corrected q < .05. **D.** Purple clusters show searchlight results of significant decoding for hand vs. eye information in contralateral (left) auditory cortex during the Delay epoch. As in Experiment 1, searchlight analyses were restricted to a mask defined by significant voxels in a Intact Speech > Rest contrast using the independent auditory localizer task data (gray traced regions; see Methods). Group-level searchlight maps were thresholded at t > 2.65 (one-tailed p < .01) and cluster-corrected at p < .05. **E.** Significant decoding in the cerebellum, for the same analysis as in D.

#### Delay period decoding from early auditory cortex is unlikely to reflect auditory rehearsal

As in Experiment 1, a pattern classification analysis on Delay period voxel patterns revealed that information related to the upcoming effector to be used (hand vs. eye) could be decoded from the STG (see Fig. 4C). Notably, we found that this decoding during the Delay epoch was only observed in the contralateral (left), and not ipsilateral (right), STG (left: t_12_ = 3.58, p = .001; right: t_12_ = 1.42, p = .091), which was also significant in follow-up permutation analyses (p = .001; see Supplemental Fig 6A). Decoding in the remaining ROIs during the Delay epoch were all p > .10. During the Execute epoch, as in Experiment 1, we found that effector-related decoding was robust in all three areas and in both hemispheres (all p < .008; all permutation analyses p < .001). Together, these results support the key observation from Experiment 1 that the auditory cortex contains effector-specific information prior to movement onset. In addition, this second experiment does not display the expected characteristics of auditory rehearsal/working memory processes (Paulesu et al. 1993; Kumar et al. 2016), in that (1) the effector-related modulation occurs in the absence of any BOLD activation prior to movement during the Delay period (Fig. 4B shows the baseline-levels of activity in both left and right STG), and that (2) this effector-specific information is represented in the auditory cortex contralateral to the (right) hand being used, rather than in bilateral auditory cortex. With respect to this latter finding, if auditory cortex is modulated in a top-down fashion via the motor system, then we might expect — given the contralateral organization of the motor system (Porter and Lemon 1995) and the existence of direct within-hemispheric projections from motor to auditory cortex (Nelson et al. 2013b) — that these modulations should be primarily contralateral (to the hand) in nature.

As in Experiment 1, it is useful to interpret these above decoding effects in left STG with respect to a positive control region, like PMd, known to distinguish upcoming hand versus eye movements during the Delay epoch in humans (Gallivan et al. 2011). As shown in the Supplementary Fig. 6B, we find a similar level of decoding of effector-specific information in left PMd (t_12_ = 2.78, p = .008) as that observed in left STG above. This shows that motor-related information is just as decodable from the auditory cortex as it is from a motor-related region, like PMd.

#### Searchlight analyses reveal the representation of effector-specific information in contralateral auditory cortex prior to movement

To complement our ROI analyses and bolster our observations in Experiment 1, we again performed a group-level searchlight analysis within the wider auditory processing network (denoted by the gray traces in Fig. 4D,E). During the Delay epoch (see Fig. 4D), we identified a large effector-specific (hand vs. eye) decoding cluster in left STG (308 voxels; peak, x = −54, y = −28, z = 4, t_13_ = 9.49, p<.00001), as well as a smaller cluster in cerebellum (150 voxels; peak, x = 6, y = −78, z = −18, t_13_ = 4.47, p<.00026, see Fig. 4E). An uncorrected searchlight map (see Supplemental Fig. 7A) also reveals no major sub-threshold clusters in right STG, reinforcing the notion that decoding in Experiment 2 is contralateral in nature. A searchlight analysis using the Execute epoch data replicated our general observations from Experiment 1 that movement execution elicits far more widespread activity through the wider auditory network (see Supplemental Fig 7). These searchlight results, when considered jointly with our ROI-based findings, provide strong evidence that movement preparation primarily modulates neural activity in the contralateral auditory cortex.

## DISCUSSION

Here we have shown, using fMRI and two separate experiments involving delayed movement tasks, that effector-specific information (i.e., left vs. right hand in Experiment 1, and hand vs. eye in Experiment 2) can be decoded from pre-movement neural activity patterns in early auditory cortex. In Experiment 1, we showed that effector-specific decoding was invariant to the auditory cue used to instruct the participant on which hand to use. Separately, with our searchlight analyses, we also found that the decoding of this hand-related information occured in a separate subregion of auditory cortex than the decoding of the auditory cue that instructed the motor action. In Experiment 2, we showed that effector-specific decoding in the auditory cortex occurs even in the absence of any auditory cues (i.e., we cued movements via a visual cue) or even when there is no net increase in BOLD activity during the Delay period. Moreover, we found through both our ROI-based and searchlight-based analyses that this decoding in auditory cortex was contralateral to the hand being used. Taken together, the findings from these two experiments suggest that a component of action planning, beyond preparing motor areas for the forthcoming movement, involves modulating early sensory cortical regions. Such modulation may enable these areas to more effectively participate in the processing of task-specific sensory signals that normally arise during the unfolding movement itself.

### ‘Motor’ versus ‘Sensory’ plans

With respect to motor-related brain areas (e.g., the primary, premotor, and supplementary motor cortices), several hypotheses have been proposed about the role of planning-related activity. Some researchers have suggested that planning activity codes a variety of different movement parameters (e.g., direction, speed), with a view that it represents some subthreshold version of the forthcoming movement (Tanji and Evarts 1976; Riehle and Requin 1989; Hocherman and Wise 1991; Shen and Alexander 1997; Messier and Kalaska 2000b; Churchland et al. 2006; Pesaran et al. 2006; Batista et al. 2007). More recent work, examining populations of neurons in motor areas, has instead suggested that movement planning involves setting up the initial state of the population, such that movement execution can unfold naturally through transitory neural dynamics (Churchland et al. 2010, 2012; Shenoy et al. 2013a; Sussillo et al. 2015; Pandarinath et al. 2017; Lara et al. 2018). In the context of this newer framework, our results suggest that motor planning may also involve setting up the initial state of primary sensory cortical areas. Whereas the neural activity patterns that unfold during movement execution in motor areas are thought to regulate the timing and nature of descending motor commands (Churchland et al. 2012; Shenoy et al. 2013b), such activity in primary sensory areas could instead regulate the timing and nature of the processing of incoming sensory signals.

One idea is that motor preparation signals originating from the motor system could tune early sensory areas for a role in sensory prediction. Prediction of the sensory consequences of action is essential for the accurate sensorimotor control of movement, per se, and also provides a mechanism for distinguishing between self-generated and externally generated sensory information (Wolpert and Flanagan 2001). The critical role of prediction in sensorimotor control has been well documented in the context of object manipulation tasks (Flanagan et al. 2006; Johansson and Flanagan 2009), like those used in our experiments. The control of such tasks centers around *contact events,* which give rise to discrete sensory signals in multiple modalities (e.g., auditory, tactile) and represent subgoals of the overall task. Thus, in the grasp, lift, and replace task that our participants performed in both Experiments 1 and 2, the brain automatically predicts the timing and nature of discrete sensory signals associated with contact between the digits and object, as well as the breaking, and subsequent making, of contact between the object and surface; events that signify successful object grasp, lift-off, and replacement, respectively. By comparing predicted and actual sensory signals associated with these events, the brain can monitor task progression and launch rapid corrective actions if mismatches occur (Wolpert et al. 2011). These corrective actions are themselves quite sophisticated and depend on both the phase of the task and the nature of the mismatch (Flanagan et al. 2006). Thus, outside the motor system, the preparation of manipulation tasks clearly involves forming what could be referred to as a *‘sensory plan’;* i.e., a series of sensory signals—linked to contact events—that, during subsequent movement execution, can be predicted based on knowledge of object properties and information related to outgoing motor commands (Johansson and Flanagan, 2009).

In the context of the experimental tasks and results presented here, it would make sense that such ‘sensory plans’ be directly linked to the acting limb (and thus, decodable at the level of auditory cortex). This is because successful sensory prediction, error detection, and rapid motor correction, would necessarily require direct knowledge of the hand being used. There is evidence from both songbirds and rodents to suggest that the internal motor-based estimate of auditory consequences is established in auditory cortex (Canopoli et al. 2014; Schneider et al. 2018). This provides a putative mechanism through which auditory cortex itself can generate sensory error signals that can be used to update ongoing movement. However, for the latter to occur, auditory cortex must be able to relay this sensory error information back to the motor system, either through direct connections, or through some intermediary brain area. In principle, this could be achieved through known projections from auditory to frontal cortical regions. Neuroanatomical tracing studies in nonhuman primates have identified bidirectional projections between regions in auditory and frontal cortex (Petrides and Pandya 1988, 2002; Deacon 1992; Romanski et al. 1999), and in humans, the arcuate and uncinated fasciculi fiber tracts are presumed to allow the bidirectional exchange of information between the auditory and motor cortices (Schneider and Mooney 2018a). However, any auditory-to-motor flow of information exchange must account for the fact that the motor system, unlike the auditory system, has a contralateral organization (Porter and Lemon 1995). That is, given that the auditory sensory reafference associated with movements of a single limb (e.g., right hand) is processed in bilateral auditory cortex, then there must be a mechanism by which resulting sensory auditory errors can be used to selectively update the contralateral motor (e.g., left) cortex.

One possibility, supported by our finding that effector-related information could only be decoded from the the contralateral auditory cortex (in Experiment 2), is that the motor hemisphere involved in movement planning (i.e., contralateral to the limb) may only exert a top-down modulation on the auditory cortex within the same hemisphere. This would not only be consistent with current knowledge in rodents that motor-to-auditory projections are within hemisphere only (Nelson et al. 2013b), but it would also provide a natural cortical mechanism by which sensory errors computed in auditory cortex can be directly tagged to the specific motor effector being used. For example, if the motor cortex contralateral to the acting hand is providing an internal motor-based estimate of the auditory consequences of movement to the auditory cortex within the same hemisphere, then only *that* auditory cortex would be able to compute the sensory error signal. In this way, the intrinsic within-hemispheric wiring of motor-to-auditory connections could provide the key architecture through which auditory error signals can be linked to (and processed with respect to) the acting hand and thus also potentially used to update motor commands directly.

One addendum that should be added to our above speculation is that, due to the loud MRI scanner environment, subjects would have *never* expected to hear any of the auditory consequences associated with their movements. On the one hand, this bolsters the view that the modulations we describe in auditory cortex are ‘automatic’ in nature (i.e., they are context-invariant and occur outside the purview of cognitive control). On the other hand, it might instead suggest that, rather than their being an interaction between hand-dependency and expected auditory consequences (as speculated in the previous paragraph), there is an effector-dependent global gating mechanism associated with movement planning. Moreover, given that in Experiment 2 we show that the effector decoding occurs in contralateral auditory cortex during planning whereas it occurs in bilateral auditory cortex during execution (Figure S7), it could suggest different gating mechanisms for action planning versus execution. Further studies will be needed to address these possibilities.

### Sensory predictions for sensory cancellation versus sensorimotor control

The disambiguation of self- and externally generated sensory information is thought to rely on cancelling, or attenuating, the predictable sensory consequences of movements (Wolpert and Flanagan 2001; Schneider and Mooney 2018b). Such sensory predictions for use in *sensory cancellation* are generally thought to be represented in primary sensory areas. According to this view, an efference copy of descending motor commands, associated with movement execution, is transmitted in a top-down fashion to early sensory cortices in order to attenuate self-generated sensory information (Holst et al. 1950; Crapse and Sommer 2008). By contrast, sensory predictions for use in *sensorimotor control* are thought to be represented in the same frontoparietal circuits involved in movement planning and execution (Scott 2012, 2016). According to this view, incoming sensory information, associated with movement execution, is transmitted in a bottom-up fashion from early sensory areas to frontoparietal circuits wherein processes associated with mismatch detection and subsequent movement correction are performed (Desmurget et al. 1999; Tunik et al. 2005; Jenmalm et al. 2006). Although here we observe our task-related modulations in early sensory (auditory) cortex, we think it is unlikely that it purely reflects a sensory cancellation process per se. First, sensory attenuation responses in primary auditory cortex have been shown to occur about two hundred *milliseconds* prior to movement onset (Schneider et al. 2014, 2018), whereas the effector-specific modulations of auditory cortex we report here occurred, at minimum, several *seconds* prior to movement onset (as revealed through our analyses linked to the onset of the Delay epoch). Second, as already noted above, any sensory attenuation would be expected to occur in *bilateral* auditory cortex (Schneider et al. 2014), which contrasts with our finding of effector-related decoding in only the contralateral auditory cortex in Experiment 2. Given these considerations, a more likely explanation of our results is that they reflect the motor system preparing the state of auditory cortex, in a top-down fashion, to process future auditory signals for a role in forthcoming *sensorimotor control.*

### Effector-specific representations in auditory cortex are likely motor-related in nature

There are three lines of support for the notion that the effector-specific decoding observed here in auditory cortex is the result of a top-down, motor-related modulation. First, as noted above, in the MRI scanner environment (i.e., loud background noise, subjects’ wearing of headphones), our participants could not have heard the auditory consequences of their actions; e.g., sounds associated with contacting, lifting, and replacing the object. This argues that our reported modulation of auditory cortex is automatic in nature, and not linked to any sensory reafference or attentional processes (see Otazu et al. 2009; Schneider et al. 2014). Second, our searchlight analyses in Experiment 1 identified separate clusters in STG that decoded hand information (in early auditory cortex) from those that decoded auditory cue information (in higher-order auditory cortex). The fact that different subregions of auditory cortex are modulated prior to movement for motor (i.e., discriminating left vs. right hand) versus auditory language information (i.e., discriminating ‘Compty’ vs. ‘Midwig’) is consistent with the notion that our hand-related decoding effects are invariant to sensory input. This finding, along with others presented in our paper (see Results), also argues against the idea that our hand-related decoding results can be attributed to some subvocalization process, as these would be expected to recruit the same region of auditory cortex as when actually hearing the commands (Paulesu et al. 1993; McGuire et al. 1996; Shergill et al. 2001). Third, in Experiment 2 the decoding of effector-specific information was found to be contralateral to the hand being used in the task. As already noted, such lateralization is a main organizational feature of movement planning and control throughout the motor system (Porter and Lemon 1995), and is consistent with neuroanatomical tracing work showing that the motor cortex has direct projections to the auditory cortex within the same hemisphere (Nelson et al. 2013b). By contrast, no such organization exists in the cortical auditory system, with lateralization instead thought to occur along different dimensions (e.g., language processing (Hackett 2015)). Notably, the contralaterality observed in our Experiment 2 also similarly rules out the possibility that our results are merely the byproduct of an auditory working-memory/rehearsal process, as these have been shown to recruit *bilateral* auditory cortex (Kumar et al. 2016). Indeed, we find through both our ROI-based and searchlight-based analyses that the decoding of motor effector information occurs in *bilateral* auditory cortex when *both* hands are interchangeably used in the task (Experiment 1) and find only contralateral decoding of motor effector information when only one of the hands is used in the task (Experiment 2). These observations, when taken together, support the notion that the signals being decoded from the auditory cortex prior to movement have a motor, rather than sensory, origin. We appreciate, however, that we are only able to infer that the modulation in auditory cortex arises via the motor system, as in the current study we do not assess any circuit-level mechanisms related to directionality or causality, as has been done in non-human work (Nelson et al. 2013a; Schneider et al. 2014).

### Representation of predicted tactile input in auditory cortex?

Prior work has demonstrated that tactile input alone is capable of driving auditory cortex activity (Foxe et al. 2002; Kayser et al. 2005; Schürmann et al. 2006; Lakatos et al. 2007), indicating a potential role for auditory cortex in multisensory integration. As noted above, the control of object manipulation tasks involves accurately predicting discrete sensory events that arise in multiple modalities, including tactile and auditory (Johansson and Flanagan 2009). It is plausible then that some of the pre-movement auditory cortex modulation described here reflects the predicted *tactile* events arising from our task (e.g., object contact, lift-off and replacement), which we would also expect to be linked to the acting hand (and thus, decodable). Though we cannot disentangle this possibility in the current study, it does not undercut our main observation that early auditory cortex is selectively modulated as a function of the movement being prepared; nor does it undercut our interpretation—that such modulation is likely linked to sensorimotor prediction.

### Conclusions

Here we show that, prior to movement, neural activity patterns in early auditory cortex carry information about the movement effector to be used in the upcoming action. This result supports the hypothesis that sensorimotor planning, which is critical in preparing neural states ahead of movement execution (Lara et al. 2018), not only occurs in motor areas but also in primary sensory areas. While further work is required to establish the precise role of this movement-related modulation, our findings add to a growing line of evidence indicating that early sensory systems are directly modulated by sensorimotor computations performed in higher-order cortex (Chapman et al. 2011; Steinmetz and Moore 2014; Gutteling et al. 2015; Gallivan et al. 2019b) and not merely low-level relayers of incoming sensory information (Scheich et al. 2007; Matyas et al. 2010; Weinberger 2011; Huang et al. 2019).

## Acknowledgements

This work was supported by operating grants from the Canadian Institutes of Health Research (CIHR) awarded to J.R.F. and J.P.G. (MOP126158). J.P.G. was supported by a Natural Sciences and Engineering Research Council (NSERC) Discovery Grant, as well as funding from the Canadian Foundation for Innovation. D.J.G. was supported by a R.S. McLaughlin Fellowship and an NSERC graduate fellowship; C.N.A. was supported by an NSERC graduate fellowship. The authors would like to thank Martin York, Sean Hickman, and Don O’Brien for technical assistance.

## Author Contributions

J.R.F., I.S.J and J.P.G. designed the experiments. D.J.G. and J.P.G. performed research. D.J.G., C.N.A., C.H., J.R.F and J.P.G., analyzed and interpreted data. D.J.G., J.R.F and J.P.G wrote the paper. All authors provided edits and feedback on the final version of the paper.

## Competing Interests Statement

The authors declare no competing financial interests.

## Notes

### Competing Interest Statement

The authors have declared no competing interest.

### Summary of Updates

Final edits and analyses are now included

## REFERENCES

Abraham A, Pedregosa F, Eickenberg M, Gervais P, Mueller A, Kossaifi J, Gramfort A, Thirion B, Varoquaux G. 2014. Machine learning for neuroimaging with scikit-learn. Front Neuroinform. 8:14.

Avants BB, Epstein CL, Grossman M, Gee JC. 2008. Symmetric diffeomorphic image registration with cross-correlation: evaluating automated labeling of elderly and neurodegenerative brain. Med Image Anal. 12:26–41.

Batista AP, Santhanam G, Yu BM, Ryu SI, Afshar A, Shenoy KV. 2007. Reference Frames for Reach Planning in Macaque Dorsal Premotor Cortex. Journal of Neurophysiology.

Canopoli A, Herbst JA, Hahnloser RHR. 2014. A higher sensory brain region is involved in reversing reinforcement-induced vocal changes in a songbird. J Neurosci. 34:7018–7026.

Chapman CS, Gallivan JP, Culham JC, Goodale MA. 2011. Mental blocks: fMRI reveals top-down modulation of early visual cortex when obstacles interfere with grasp planning. Neuropsychologia. 49:1703–1717.

Churchland MM, Cunningham JP, Kaufman MT, Foster JD, Nuyujukian P, Ryu SI, Shenoy KV. 2012. Neural population dynamics during reaching. Nature. 487:51–56.

Churchland MM, Cunningham JP, Kaufman MT, Ryu SI, Shenoy KV. 2010. Cortical preparatory activity: representation of movement or first cog in a dynamical machine? Neuron. 68:387–400.

Churchland MM, Santhanam G, Shenoy KV. 2006. Preparatory activity in premotor and motor cortex reflects the speed of the upcoming reach. J Neurophysiol. 96:3130–3146.

Cisek P, Crammond DJ, Kalaska JF. 2003. Neural activity in primary motor and dorsal premotor cortex in reaching tasks with the contralateral versus ipsilateral arm. J Neurophysiol. 89:922–942.

Cox RW. 1996. AFNI: software for analysis and visualization of functional magnetic resonance neuroimages. Comput Biomed Res. 29:162–173.

Cox RW, Chen G, Glen DR, Reynolds RC, Taylor PA. 2017. FMRI Clustering in AFNI: False-Positive Rates Redux. Brain Connect. 7:152–171.

Cox RW, Hyde JS. 1997. Software tools for analysis and visualization of fMRI data. NMR Biomed. 10:171–178.

Crapse TB, Sommer MA. 2008. Corollary discharge across the animal kingdom. Nat Rev Neurosci. 9:587–600.

Cui H, Andersen RA. 2007. Posterior parietal cortex encodes autonomously selected motor plans. Neuron. 56:552–559.

Da Costa S, van der Zwaag W, Marques JP, Frackowiak RSJ, Clarke S, Saenz M. 2011. Human primary auditory cortex follows the shape of Heschl’s gyrus. J Neurosci. 31:14067–14075.

Dale AM, Fischl B, Sereno MI. 1999. Cortical surface-based analysis. I. Segmentation and surface reconstruction. Neuroimage. 9:179–194.

Deacon TW. 1992. Cortical connections of the inferior arcuate sulcus cortex in the macaque brain. Brain Res. 573:8–26.

Desikan RS, Ségonne F, Fischl B, Quinn BT, Dickerson BC, Blacker D, Buckner RL, Dale AM, Maguire RP, Hyman BT, Albert MS, Killiany RJ. 2006. An automated labeling system for subdividing the human cerebral cortex on MRI scans into gyral based regions of interest. Neuroimage. 31:968–980.

Desmurget M, Epstein CM, Turner RS, Prablanc C, Alexander GE, Grafton ST. 1999. Role of the posterior parietal cortex in updating reaching movements to a visual target. Nat Neurosci. 2:563–567.

Eliades SJ, Wang X. 2008. Neural substrates of vocalization feedback monitoring in primate auditory cortex. Nature.

Esteban O, Markiewicz CJ, Blair RW, Moodie CA, Ilkay Isik A, Erramuzpe A, Kent JD, Goncalves M, DuPre E, Snyder M, Oya H, Ghosh SS, Wright J, Durnez J, Poldrack RA, Gorgolewski KJ. 2018. fMRIPrep: a robust preprocessing pipeline for functional MRI. Nat Methods. 1.

Esteban O, Markiewicz CJ, Burns C, Johnson H, Ziegler E, Manhães-Savio A, Jarecka D, Ellis DG, Yvernault B, Hamalainen C, Notter MP, Salo T, Waskom M, Goncalves M, Jordan K, Wong J, Dewey BE, Madison C, Clark D, Loney F, Clark D, Nielson DM, Keshavan A, Joseph M, Dayan M, Modat M, Bougacha S, Gramfort A, Visconti di Oleggio Castello M, Pinsard B, Berleant S, Christian H, Rokem A, Halchenko YO, Kaczmarzyk J, Benderoff E, Ćirić R, Varoquaux G, Moloney B, DuPre E, Koudoro S, Clark MG, Wassermann D, Cipollini B, Guillon J, Markello R, Buchanan C, Hanke M, Tungaraza R, Sikka S, Gillman A, Pauli WM, de Hollander G, Forbes J, Iqbal S, Mordom D, Mancini M, Malone IB, Dubois M, Schwartz Y, Frohlich C, Tabas A, Welch D, Richie-Halford A, Tilley S II, Watanabe A, Nichols BN, Huntenburg JM, Eshaghi A, Schaefer A, Ginsburg D, Bottenhorn K, Cumba C, Acland B, Heinsfeld AS, Kastman E, Kent J, Kleesiek J, Erickson D, Giavasis S, Ghayoor A, Liem F, De La Vega A, Küttner R, Millman J, Perez-Guevara MF, Lee JA, Zhou D, Haselgrove C, Glen D, Renfro M, Correa C, Liu S, Lampe L, Kong X-Z, Hallquist M, Kahn AE, Glatard T, Triplett W, Chawla K, Salvatore J, Pérez-García F, Ma F, Park A, Craddock RC, Hinds O, Poldrack R, Perkins LN, Kim S, Chetverikov A, Inati S, Cieslak M, Grignard M, Snoek L, Sisk LM, Leinweber K, Junhao WEN, Matsubara K, Urchs S, Blair R, Floren A, Mattfeld A, Gerhard S, Cooper G, Haehn D, Tambini A, Broderick W, Andberg SK, Noel M, Durnez J, Stadler J, Condamine E, Papadopoulos Orfanos D, Geisler D, Weinstein A, Harms R, Khanuja R, Sharp P, Stanley O, Lee N, Crusoe MR, Brett M, Falkiewicz M, Podranski K, Linkersdörfer J, Flandin G, Lerma-Usabiaga G, Shachnev D, Tarbert C, Cheung B, Meyers B, Van A, Davison A, Weninger L, Molina-Romero M, Rothmei S, Bilgel M, Schlamp K, Ort E, McNamee D, Lai J, Arias J, Bielievtsov D, Steele CJ, Huang L, Gonzalez I, Warner J, Margulies DS, Contier O, Marina A, Saase V, Nickson T, Varada J, Schwabacher I, Pellman J, Khanuja R, Pannetier N, McDermottroe C, Mihai PG, Lai J, Gorgolewski KJ, Ghosh S. 2019. Nipy/nipype: 1.4.0.

Evans AC, Janke AL, Collins DL, Baillet S. 2012. Brain templates and atlases. Neuroimage. 62:911–922.

Flanagan JR, Bowman MC, Johansson RS. 2006. Control strategies in object manipulation tasks. Curr Opin Neurobiol. 16:650–659.

Foxe JJ, Wylie GR, Martinez A, Schroeder CE, Javitt DC, Guilfoyle D, Ritter W, Murray MM. 2002. Auditory-somatosensory multisensory processing in auditory association cortex: an fMRI study. J Neurophysiol. 88:540–543.

Gallivan JP, Cant JS, Goodale MA, Flanagan JR. 2014. Representation of object weight in human ventral visual cortex. Curr Biol. 24:1866–1873.

Gallivan JP, Chapman CS, Gale DJ, Flanagan JR, Culham JC. 2019a. Selective Modulation of Early Visual Cortical Activity by Movement Intention. Cereb Cortex.

Gallivan JP, Chapman CS, Gale DJ, Flanagan JR, Culham JC. 2019b. Selective Modulation of Early Visual Cortical Activity by Movement Intention. Cereb Cortex.

Gallivan JP, Chapman CS, McLean DA, Flanagan JR, Culham JC. 2013. Activity patterns in the category-selective occipitotemporal cortex predict upcoming motor actions. Eur J Neurosci. 38:2408–2424.

Gallivan JP, Johnsrude IS, Flanagan JR. 2016. Planning Ahead: Object-Directed Sequential Actions Decoded from Human Frontoparietal and Occipitotemporal Networks. Cereb Cortex. 26:708–730.

Gallivan JP, McLean DA, Flanagan JR, Culham JC. 2013. Where one hand meets the other: limb-specific and action-dependent movement plans decoded from preparatory signals in single human frontoparietal brain areas. J Neurosci. 33:1991–2008.

Gallivan JP, McLean DA, Smith FW, Culham JC. 2011. Decoding effector-dependent and effector-independent movement intentions from human parieto-frontal brain activity. J Neurosci. 31:17149–17168.

Gallivan JP, McLean DA, Valyear KF, Culham JC. 2013. Decoding the neural mechanisms of human tool use. Elife. 2:e00425.

Gorgolewski K, Burns CD, Madison C, Clark D, Halchenko YO, Waskom ML, Ghosh SS. 2011. Nipype: a flexible, lightweight and extensible neuroimaging data processing framework in python. Front Neuroinform. 5:13.

Greve DN, Fischl B. 2009. Accurate and robust brain image alignment using boundary-based registration. Neuroimage. 48:63–72.

Gutteling TP, Petridou N, Dumoulin SO, Harvey BM, Aarnoutse EJ, Kenemans JL, Neggers SFW. 2015. Action preparation shapes processing in early visual cortex. J Neurosci. 35:6472–6480.

Hackett TA. 2015. Anatomic organization of the auditory cortex. Handb Clin Neurol. 129:27–53.

Hackett TA, de la Mothe LA, Camalier CR, Falchier A, Lakatos P, Kajikawa Y, Schroeder CE. 2014. Feedforward and feedback projections of caudal belt and parabelt areas of auditory cortex: refining the hierarchical model. Front Neurosci. 8:72.

Hackett TA, Preuss TM, Kaas JH. 2001. Architectonic identification of the core region in auditory cortex of macaques, chimpanzees, and humans. J Comp Neurol. 441:197–222.

Hocherman S, Wise SP. 1991. Effects of hand movement path on motor cortical activity in awake, behaving rhesus monkeys. Exp Brain Res. 83:285–302.

Holst E von, von Holst E, Mittelstaedt H. 1950. Das Reafferenzprinzip. Naturwissenschaften.

Huang Y, Heil P, Brosch M. 2019. Associations between sounds and actions in early auditory cortex of nonhuman primates. Elife. 8.

Hutchison RM, Gallivan JP. 2018. Functional coupling between frontoparietal and occipitotemporal pathways during action and perception. Cortex. 98:8–27.

Jenkinson M, Bannister P, Brady M, Smith S. 2002. Improved optimization for the robust and accurate linear registration and motion correction of brain images. Neuroimage. 17:825–841.

Jenmalm P, Schmitz C, Forssberg H, Ehrsson HH. 2006. Lighter or heavier than predicted: neural correlates of corrective mechanisms during erroneously programmed lifts. J Neurosci. 26:9015–9021.

Johansson, Flanagan JR. 2009. Coding and use of tactile signals from the fingertips in object manipulation tasks. Nat Rev Neurosci. 10:345–359.

Kaufman MT, Churchland MM, Santhanam G, Yu BM, Afshar A, Ryu SI, Shenoy KV. 2010. Roles of monkey premotor neuron classes in movement preparation and execution. J Neurophysiol. 104:799–810.

Kayser C, Petkov CI, Augath M, Logothetis NK. 2005. Integration of touch and sound in auditory cortex. Neuron. 48:373–384.

Keck T, Keller GB, Jacobsen RI, Eysel UT, Bonhoeffer T, Hübener M. 2013. Synaptic scaling and homeostatic plasticity in the mouse visual cortex in vivo. Neuron. 80:327–334.

Keller GB, Bonhoeffer T, Hübener M. 2012. Sensorimotor mismatch signals in primary visual cortex of the behaving mouse. Neuron. 74:809–815.

Klein A, Ghosh SS, Bao FS, Giard J, Häme Y, Stavsky E, Lee N, Rossa B, Reuter M, Chaibub Neto E, Keshavan A. 2017. Mindboggling morphometry of human brains. PLoS Comput Biol. 13:e1005350.

Kriegeskorte N, Goebel R, Bandettini P. 2006. Information-based functional brain mapping. Proc Natl Acad Sci U S A. 103:3863–3868.

Kumar S, Joseph S, Gander PE, Barascud N, Halpern AR, Griffiths TD. 2016. A Brain System for Auditory Working Memory. Journal of Neuroscience.

Lakatos P, Chen C-M, O’Connell MN, Mills A, Schroeder CE. 2007. Neuronal oscillations and multisensory interaction in primary auditory cortex. Neuron. 53:279–292.

Lanczos C. 1964. Evaluation of Noisy Data. Journal of the Society for Industrial and Applied Mathematics Series B Numerical Analysis. 1:76–85.

Lara, Elsayed GF, Zimnik AJ, Cunningham JP, Churchland MM. 2018. Conservation of preparatory neural events in monkey motor cortex regardless of how movement is initiated. Elife. 7.

Lee AM, Moses Lee A, Hoy JL, Bonci A, Wilbrecht L, Stryker MP, Niell CM. 2014. Identification of a Brainstem Circuit Regulating Visual Cortical State in Parallel with Locomotion. Neuron.

Leinweber M, Ward DR, Sobczak JM, Attinger A, Keller GB. 2017. A Sensorimotor Circuit in Mouse Cortex for Visual Flow Predictions. Neuron. 96:1204.

Linke AC, Cusack R. 2015. Flexible information coding in human auditory cortex during perception, imagery, and STM of complex sounds. J Cogn Neurosci. 27:1322–1333.

Mandelblat-Cerf Y, Las L, Denisenko N, Fee MS. 2014. A role for descending auditory cortical projections in songbird vocal learning. Elife. 3.

Matyas F, Sreenivasan V, Marbach F, Wacongne C, Barsy B, Mateo C, Aronoff R, Petersen CCH. 2010. Motor control by sensory cortex. Science. 330:1240–1243.

McGuire PK, Silbersweig DA, Frith CD. 1996. Functional neuroanatomy of verbal self-monitoring. Brain. 119 (Pt 3):907–917.

Messier J, Kalaska JF. 2000a. Covariation of primate dorsal premotor cell activity with direction and amplitude during a memorized-delay reaching task. J Neurophysiol. 84:152–165.

Messier J, Kalaska JF. 2000b. Covariation of primate dorsal premotor cell activity with direction and amplitude during a memorized-delay reaching task. J Neurophysiol. 84:152–165.

Morosan P, Mohlberg H, Amunts K, Schleicher A, Rademacher J, Schormann T, Zilles K. 2000. Population maps of cytoarchitectonically defined human auditory areas. NeuroImage.

Morosan P, Rademacher J, Schleicher A, Amunts K, Schormann T, Zilles K. 2001. Human primary auditory cortex: cytoarchitectonic subdivisions and mapping into a spatial reference system. Neuroimage. 13:684–701.

Mumford JA, Turner BO, Ashby FG, Poldrack RA. 2012. Deconvolving BOLD activation in event-related designs for multivoxel pattern classification analyses. Neuroimage. 59:2636–2643.

Nelson A, Schneider DM, Takatoh J, Sakurai K, Wang F, Mooney R. 2013a. A circuit for motor cortical modulation of auditory cortical activity. J Neurosci. 33:14342–14353.

Nelson A, Schneider DM, Takatoh J, Sakurai K, Wang F, Mooney R. 2013b. A Circuit for Motor Cortical Modulation of Auditory Cortical Activity. Journal of Neuroscience.

Oldfield RC. 1971. The assessment and analysis of handedness: the Edinburgh inventory. Neuropsychologia. 9:97–113.

Otazu GH, Tai L-H, Yang Y, Zador AM. 2009. Engaging in an auditory task suppresses responses in auditory cortex. Nat Neurosci. 12:646–654.

Pandarinath C, Nuyujukian P, Blabe CH, Sorice BL, Saab J, Willett FR, Hochberg LR, Shenoy KV, Henderson JM. 2017. High performance communication by people with paralysis using an intracortical brain-computer interface. Elife. 6.

Paulesu E, Frith CD, Frackowiak RS. 1993. The neural correlates of the verbal component of working memory. Nature. 362:342–345.

Pesaran B, Nelson MJ, Andersen RA. 2006. Dorsal premotor neurons encode the relative position of the hand, eye, and goal during reach planning. Neuron. 51:125–134.

Petrides M, Pandya DN. 1988. Association fiber pathways to the frontal cortex from the superior temporal region in the rhesus monkey. J Comp Neurol. 273:52–66.

Petrides M, Pandya DN. 2002. Comparative cytoarchitectonic analysis of the human and the macaque ventrolateral prefrontal cortex and corticocortical connection patterns in the monkey. Eur J Neurosci. 16:291–310.

Poeppel D, Overath T, Popper AN, Fay RR. 2012. The Human Auditory Cortex. Springer Science & Business Media.

Porter R, Lemon R. 1995. Corticospinal Function and Voluntary Movement. Clarendon Press.

Poulet JFA, Hedwig B. 2002. A corollary discharge maintains auditory sensitivity during sound production. Nature. 418:872–876.

Rademacher J, Morosan P, Schormann T, Schleicher A, Werner C, Freund HJ, Zilles K. 2001. Probabilistic mapping and volume measurement of human primary auditory cortex. Neuroimage. 13:669–683.

Ren J, Liu H, Xu T, Wang D, Li M, Lin Y, Ramirez JSB, Lu J, Li L, Ahveninen J. 2020. Individual variability in functional organization of the human and monkey auditory cortex. bioRxiv.

Reznik D, Henkin Y, Schadel N, Mukamel R. 2014. Lateralized enhancement of auditory cortex activity and increased sensitivity to self-generated sounds. Nat Commun. 5:4059.

Reznik D, Ossmy O, Mukamel R. 2015. Enhanced auditory evoked activity to self-generated sounds is mediated by primary and supplementary motor cortices. J Neurosci. 35:2173–2180.

Riehle A, Requin J. 1989. Monkey primary motor and premotor cortex: single-cell activity related to prior information about direction and extent of an intended movement. J Neurophysiol. 61:534–549.

Romanski LM, Bates JF, Goldman-Rakic PS. 1999. Auditory belt and parabelt projections to the prefrontal cortex in the rhesus monkey. J Comp Neurol. 403:141–157.

Safstrom D, Johansson RS, Flanagan JR. 2014. Gaze behavior when learning to link sequential action phases in a manual task. Journal of Vision.

Saleem AB, Ayaz A, Jeffery KJ, Harris KD, Carandini M. 2013. Integration of visual motion and locomotion in mouse visual cortex. Nat Neurosci. 16:1864–1869.

Scheich H, Brechmann A, Brosch M, Budinger E, Ohl FW. 2007. The cognitive auditory cortex: task-specificity of stimulus representations. Hear Res. 229:213–224.

Schneider DM, Mooney R. 2018a. How Movement Modulates Hearing. Annual Review of Neuroscience.

Schneider DM, Mooney R. 2018b. How Movement Modulates Hearing. Annu Rev Neurosci. 41:553–572.

Schneider DM, Nelson A, Mooney R. 2014. A synaptic and circuit basis for corollary discharge in the auditory cortex. Nature. 513:189–194.

Schneider DM, Sundararajan J, Mooney R. 2018. A cortical filter that learns to suppress the acoustic consequences of movement. Nature. 561:391–395.

Schürmann M, Caetano G, Hlushchuk Y, Jousmäki V, Hari R. 2006. Touch activates human auditory cortex. Neuroimage. 30:1325–1331.

Scott SH. 2012. The computational and neural basis of voluntary motor control and planning. Trends Cogn Sci. 16:541–549.

Scott SH. 2016. A Functional Taxonomy of Bottom-Up Sensory Feedback Processing for Motor Actions. Trends Neurosci. 39:512–526.

Shen L, Alexander GE. 1997. Preferential representation of instructed target location versus limb trajectory in dorsal premotor area. J Neurophysiol. 77:1195–1212.

Shenoy KV, Sahani M, Churchland MM. 2013a. Cortical control of arm movements: a dynamical systems perspective. Annu Rev Neurosci. 36:337–359.

Shenoy KV, Sahani M, Churchland MM. 2013b. Cortical control of arm movements: a dynamical systems perspective. Annu Rev Neurosci. 36:337–359.

Shergill SS, Bullmore ET, Brammer MJ, Williams SC, Murray RM, McGuire PK. 2001. A functional study of auditory verbal imagery. Psychol Med. 31:241–253.

Snyder LH, Batista AP, Andersen RA. 1997. Coding of intention in the posterior parietal cortex. Nature. 386:167–170.

Steinmetz NA, Moore T. 2014. Eye Movement Preparation Modulates Neuronal Responses in Area V4 When Dissociated from Attentional Demands. Neuron.

Stelzer J, Chen Y, Turner R. 2013. Statistical inference and multiple testing correction in classification-based multi-voxel pattern analysis (MVPA): random permutations and cluster size control. Neuroimage. 65:69–82.

Sussillo D, Churchland MM, Kaufman MT, Shenoy KV. 2015. A neural network that finds a naturalistic solution for the production of muscle activity. Nat Neurosci. 18:1025–1033.

Tanji J, Evarts EV. 1976. Anticipatory activity of motor cortex neurons in relation to direction of an intended movement. J Neurophysiol. 39:1062–1068.

Tunik E, Frey SH, Grafton ST. 2005. Virtual lesions of the anterior intraparietal area disrupt goal-dependent on-line adjustments of grasp. Nat Neurosci. 8:505–511.

Tustison NJ, Avants BB, Cook PA, Zheng Y, Egan A, Yushkevich PA, Gee JC. 2010. N4ITK: improved N3 bias correction. IEEE Trans Med Imaging. 29:1310–1320.

Weinberger NM. 2011. Reconceptualizing the Primary Auditory Cortex: Learning, Memory and Specific Plasticity. The Auditory Cortex.

Weinrich M, Wise SP, Mauritz KH. 1984. A neurophysiological study of the premotor cortex in the rhesus monkey. Brain. 107 (Pt 2):385–414.

Werner-Reiss U, Kelly KA, Trause AS, Underhill AM, Groh JM. 2003. Eye Position Affects Activity in Primary Auditory Cortex of Primates. Current Biology.

Wolpert DM, Diedrichsen J, Flanagan JR. 2011. Principles of sensorimotor learning. Nat Rev Neurosci. 12:739–751.

Wolpert DM, Flanagan JR. 2001. Motor prediction. Current Biology.

Wolpert DM, Miall RC. 1996. Forward Models for Physiological Motor Control. Neural Netw. 9:1265–1279.

Zhang Y, Brady M, Smith S. 2001. Segmentation of brain MR images through a hidden Markov random field model and the expectation-maximization algorithm. IEEE Trans Med Imaging. 20:45–57.

Zoellner S, Benner J, Zeidler B, Seither-Preisler A, Christiner M, Seitz A, Goebel R, Heinecke A, Wengenroth M, Blatow M, Schneider P. 2019. Reduced cortical thickness in Heschl’s gyrus as an in vivo marker for human primary auditory cortex. Hum Brain Mapp. 40:1139–1154.

