## Supplemental Data for "Motor planning modulates neural activity patterns in early human auditory cortex"

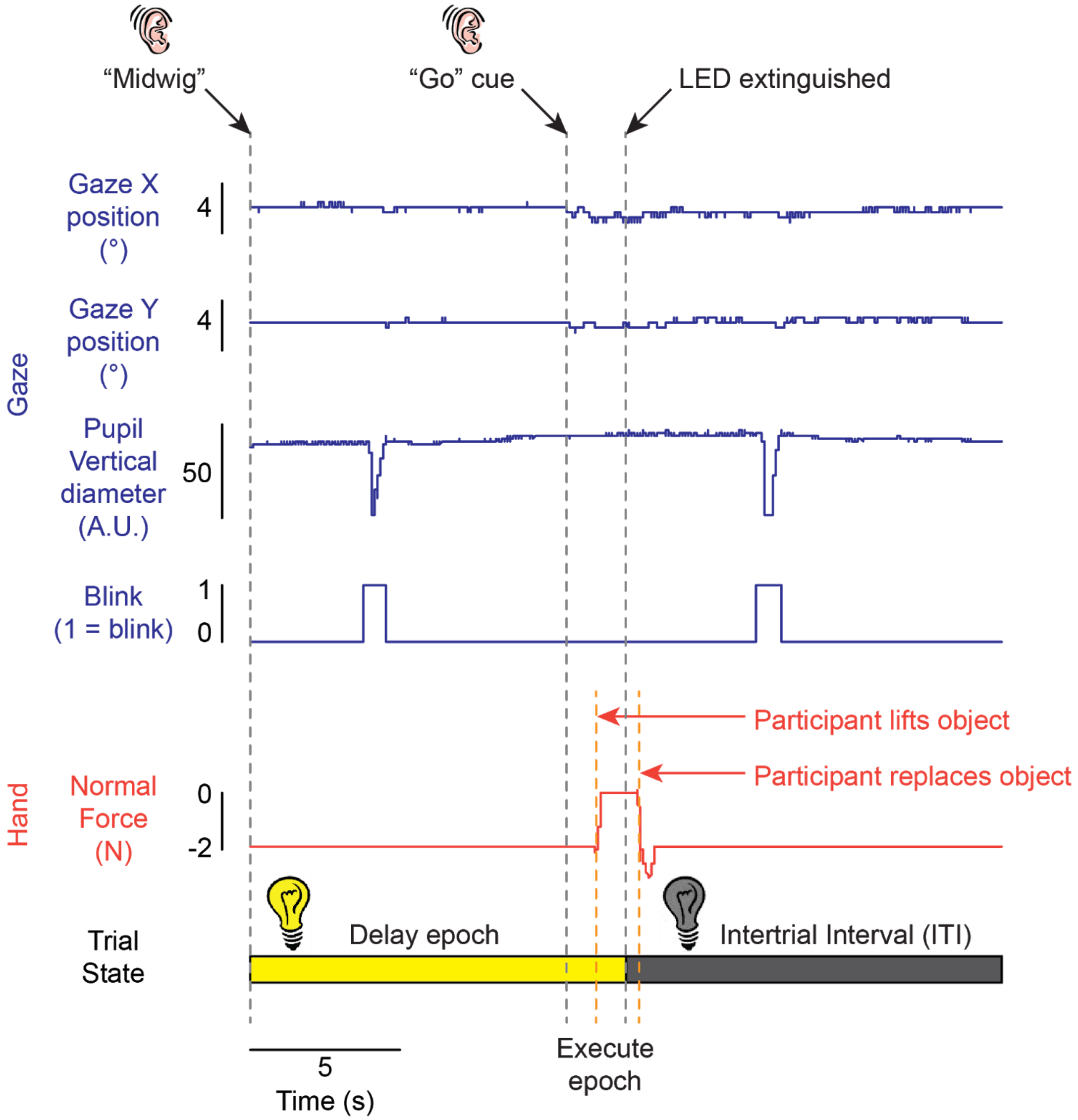


**Supplemental Figure 1. Subjects were successful at maintaining fixation during the task.** Horizontal and vertical gaze positions, pupil diameter, blink state, and recordings from force sensors located beneath the object for an example trial from a representative participant during behavioural testing for Experiment 1. Across all our gaze analyses, we observed a total of 6 saccades out of 960 total trials (16 participants x 60 trials each; 0.006% saccade rate), all of which occurred after the object was replaced (i.e., these were never made during the Delay epoch). This indicates that participants were able to maintain gaze at the fixation point, as required by the task.

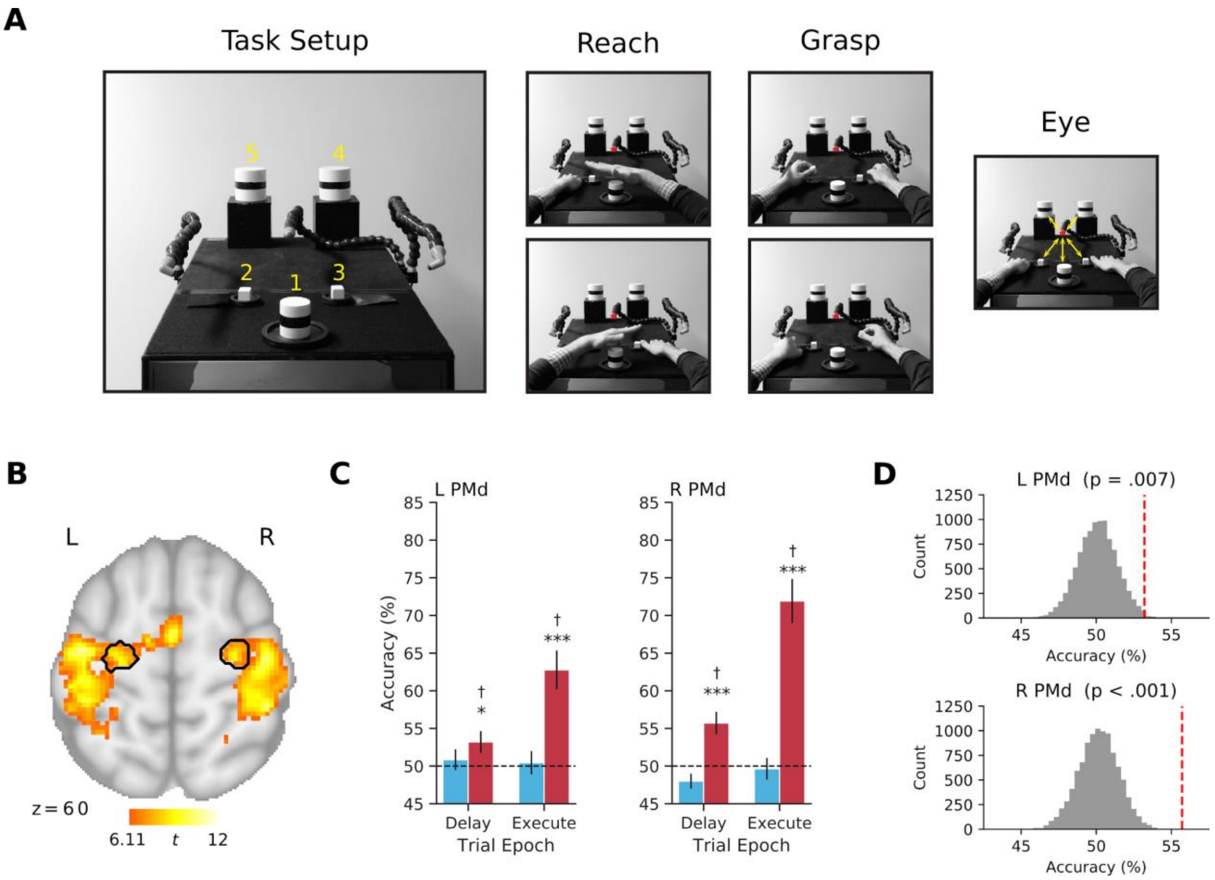


**Supplemental Figure 2. Motor localizer and dorsal premotor cortex (PMd) decoding during the main experimental task in Experiment 1**.

**A.** The motor localizer task setup and example subtasks. Eight conditions were included in this localizer, and are listed below (*note that only Grasp condition is used for analysis in this study; the remaining conditions were acquired for analysis in a separate study using the same cohort of individuals*): (1) Reach movements, which involved subjects performing alternating left and right planar limb movements (flexions about the elbow) towards the centrally located object #1 (5 x 5.5 cm, radius x height). (2) Grasp movements, which involved subjects performing alternating left and right hand precision grasp-and-lift actions (using the thumb and index finger) on small object cubes located adjacent to the start position of the hands (objects # 2 and 3, each were 2.5 × 2.5 × 2.5 cm, width × length × height). (3) Eye movements, which involved center-out-and-return eye movement fixations to five targets in the visual scene (objects #1-5; objects #4 and 5 were sized 8 x 8.5 cm, radius x height). At the beginning of each eye movement block, subjects were instructed to make their first saccade to object location #5, and then proceed in a clockwise fashion until completion of the block. (4) Finger movements, which involved subjects performing alternating left and right finger tapping (drumming) movements with their index fingers adapted from(adapted from Huth et al., 2012). (5) Touch movements, which involved subjects simultaneously rubbing, with the index fingers of their left and right hands, respectively, sandpaper or silk surfaces. Subjects were told to focus on differences in the two textures. We hypothesized that this condition would elicit maximal tactile-related activity. (6) Toe movements, which involved subjects simultaneously wiggling their left and right big toes adapted from(adapted from Huth et al., 2012). (7) Speech generation, which involved subjects performing a mental word-generation task wherein they progressed through the alphabet, silently naming a word beginning with each letter (e.g., “Ant”, “Bat”, “Car”, “Dog”, etc.) adapted from(adapted from Huth et al., 2012). (8) Mouth movements, which involved subjects repeating the word “ba-la” quietly, moving only their lips and tongue (while attempting to keep their jaw still), adapted from (adapted from Huth et al., 2012). A verbal cue, played through headphones, initiated the onset of each task block, wherein the subject heard one of the following auditory commands: “arm” (Reach), “grasp”, “eyes”, “fingers”, “touch”, “toes”, “speech”, or “mouth”. Upon cuing, subjects were then required to immediately perform the required movements (noted above) in a self-paced fashion, at an approximate rate of 1-0.5 Hz, and continue until cued otherwise at the onset of the next block. Subjects were instructed to keep the general timing of their actions as consistent as possible across trials. Other than the execution of the different hand actions, subjects were instructed to keep their hands still and in the pre-specified “home positions” on the left and right sides of the platform, and return to these same positions following movement execution. In addition, other than the eye movement condition, subjects were required to always maintain central fixation. Each trial block lasted 20 s and alternated, in a pseudo-random order, in sets of 8 and was repeated four times within an experimental run. In addition, a 20 s “rest” block, wherein subjects simply maintained central fixation, was placed at the beginning, middle and end of each run (i.e., 3 ‘rest’ blocks/run). Each experimental run totaled 700 s and subjects completed 4 of these runs during testing (resulting in a total of 16 repetitions per experimental condition per subject).

**B.** Left and right PMd (black borders). First-level contrasts of Grasp > Baseline from the motor localizer task were combined and analyzed at the group-level and thresholded at $t > 6.11$ ($p < 10^{-5}$). Then, left and right PMd were localized by centering a sphere mask (radius = 6mm) on the peak voxel located at the junction of the precentral sulcus and superior frontal sulcus (left, x = -22, y = -16, z = 60;  right, x = 28, y = -14, z = 62), and intersecting each sphere with the thresholded group contrast map. All subsequent analyses followed identical procedures to the analyses performed on the auditory regions.

**C.** Significant decoding of hand, but not cue information, in PMd during the Delay (left, $t_{14} = 2.21$, $p = .022$; right, $t_{14} = 3.84$, $p < .001$) and Execute (left, $t_{14} = 4.95$, $p < .001$; right, $t_{14} = 7.49$, $p < .001$) epochs.

**D.** The mean classification accuracies (red dashed line) for hand decoding versus their respective null distributions obtained using a two-step permutation approach (see Methods) during the Delay Epoch.

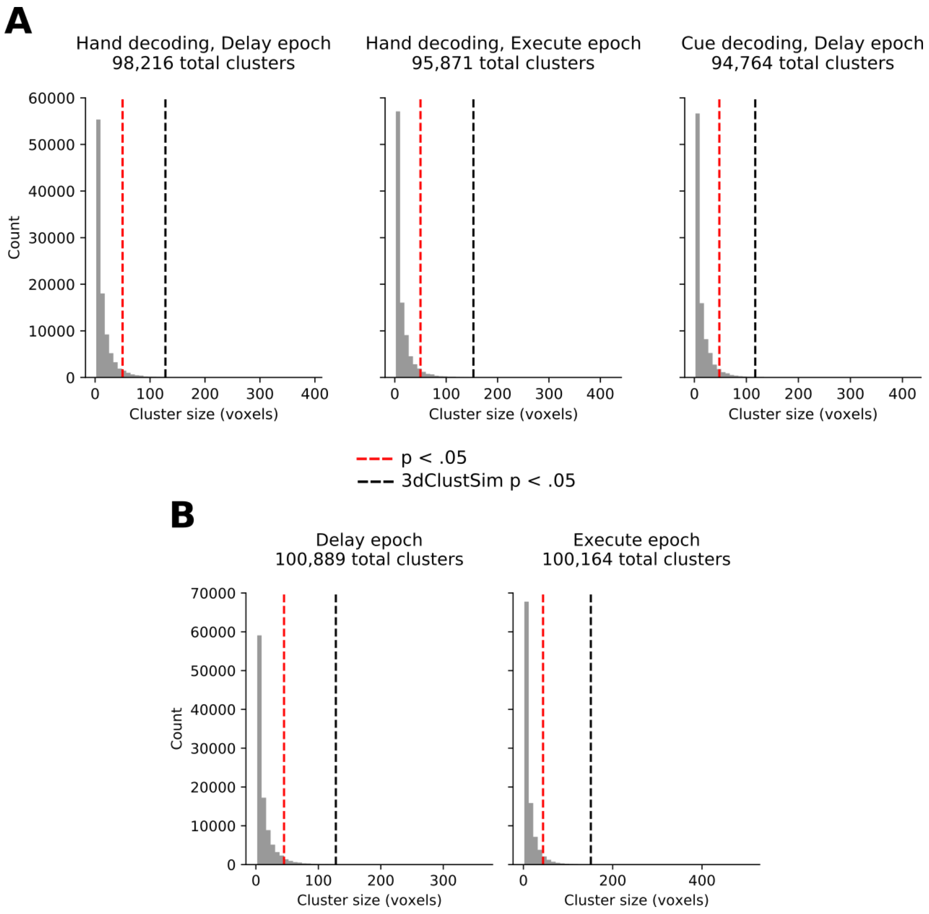


**Supplemental Figure 3. Cluster size distributions of searchlight permutation analyses for Experiment 1 and 2.** We performed an additional cluster correction approach on shuffled searchlight maps. In this approach, 100 chance decoding maps for each subject are constructed by repeatedly applying our searchlight procedure (see Methods) with randomly shuffled class labels. Then, a distribution of cluster sizes were generated by $10e^3$ iterations of a) selecting and smoothing (6mm FWHM Gaussian kernel) a random decoding map from each subject b) performing one-tailed t-tests versus 50% chance decoding on each voxel, and c) thresholding the map at $p < .01$ and extracting the sizes of all individual clusters. The cluster correction threshold was then determined by taking the minimum cluster size at which $p < .05$. Note that this approach is based on (Markiewicz & Bohland, 2016), which provides a computationally feasible alternative to (Stelzer et al., 2013) for searchlight permutation testing. **A.** Cluster size distributions for Experiment 1 searchlights. **B.** Cluster size distributions for Experiment 2 searchlights. Red dashed line indicates the determined cluster correction threshold, defined as the minimum cluster size at which $p < .05$. For comparison, cluster correction thresholds from AFNI's 3dClustSim are indicated by black dashed lines.

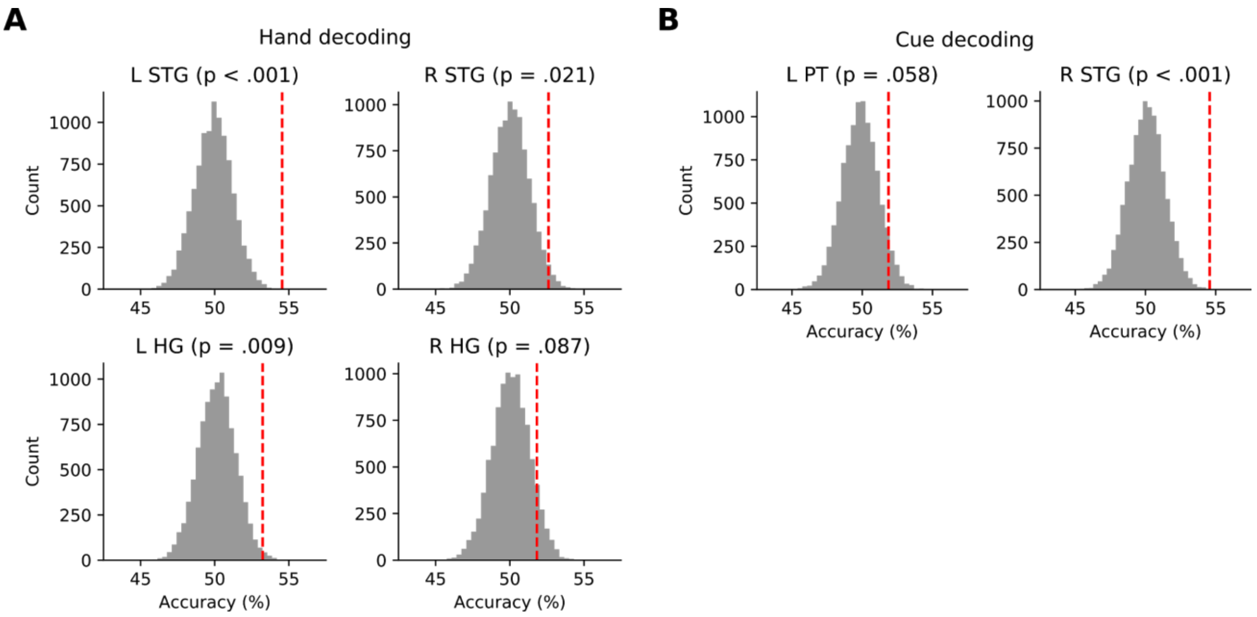


**Supplemental Figure 4. Null distribution comparisons in ROIs with significant decoding during the Delay epoch of Experiment 1. A.** Mean classification accuracy (red dashed line) versus the null distribution obtained using a two-step permutation approach (see Methods) for each ROI with significant hand decoding. **B.** Mean classification accuracy versus the null distribution in ROIs with significant cue decoding. All results shown here are consistent with the original analyses. That is, all ROIs in A and B show $p < .05$ (one-tailed), with the exception of right HG (hand decoding) and left PT (cue decoding), which did not survive correction for multiple-comparisons in the original analysis (see Figure 2).

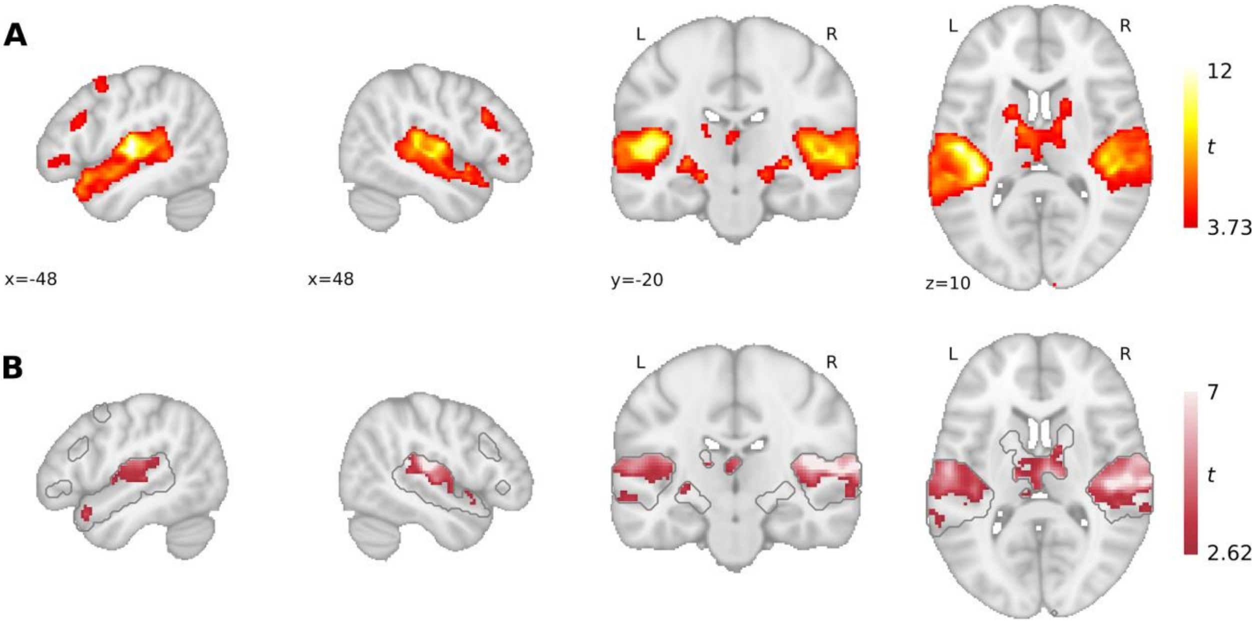


**Supplemental Figure 5. Searchlight mask and Execute epoch hand-decoding map for Experiment 1. A.** Higher-order auditory network defined at the group-level (N=16) with the contrast of Intact Speech > Rest using the independent auditory localizer task data. This network mask was used to constrain all our searchlight analyses. **B.** Cluster-corrected searchlight map for hand decoding during the Execute epoch within the boundaries of the searchlight mask. Six large clusters were identified at the following regions: right STG (2061 voxels; peak x = 44, y = -20, z = 16, $t_{14}$ = 9.40), left STG (1481 voxels; peak x = -56, y = -14, z = 14; $t_{14}$ = 6.56), left thalamus (761 voxels; peak, x = -12, y = -28, z = 2, $t_{14}$ = 9.33), vermis IV (450 voxels; peak x = -2, y = -74, z = -16, $t_{14}$ = 6.02), left middle temporal gyrus (178 voxels; peak: x = -64, y = -24, z = -8, $t_{14}$ = 5.25), and right amygdala (size = 170 voxels; peak x = 14, y = -8, z = -12, $t_{14}$ = 4.09).

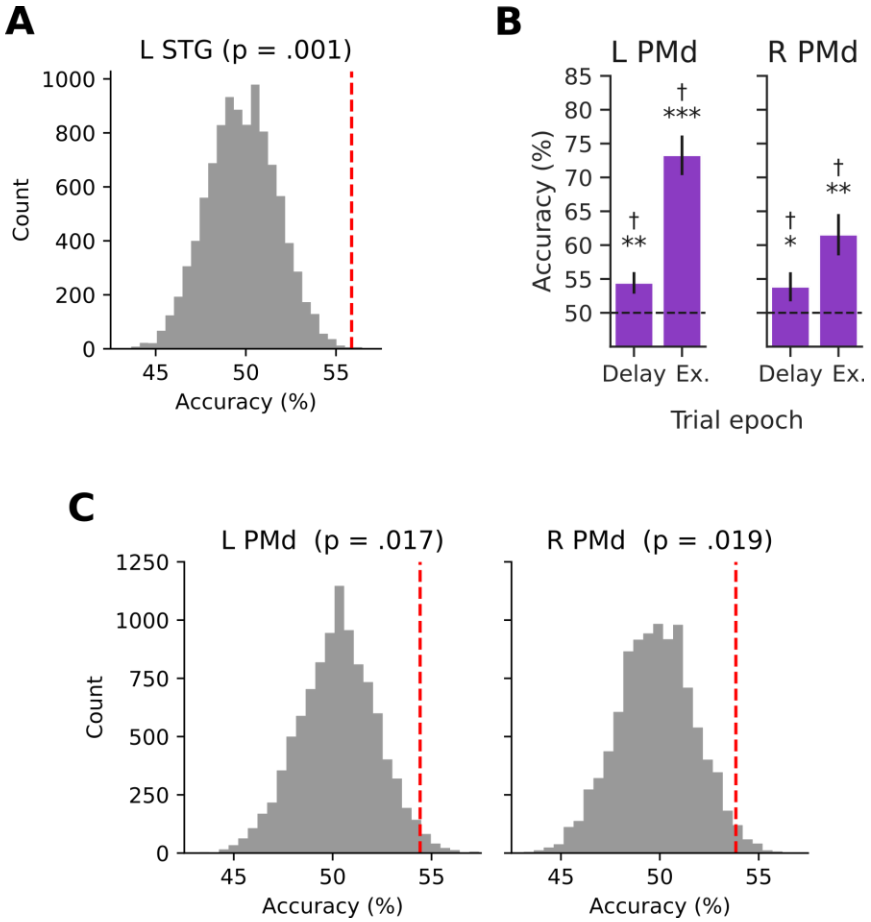


**Supplemental Figure 6. Supplemental decoding analyses for Experiment 2. A.** Comparison of the mean decoding accuracy (red dashed line) versus the null distribution obtained using a two-step permutation approach (see Methods) for left STG during the Delay epoch. **B.** Significant decoding of the effector in left and right PMd during the Delay (left, $t_{14} = 2.78$, $p = .008$; right, $t_{14} = 1.79$, $p = .049$) and Execute (Ex.; left, $t_{14} = 7.95$, $< .001$; right, $t_{14} = 3.78$, $p = .003$) epochs. Error bars show ± 1 SE of mean. * $p < .05$, ** $p < .01$, *** $p < .001$, † FDR corrected $q < .05$. **C.** Null decoding distributions in comparison with the mean decoding accuracy (red dashed line) for left and right PMd.

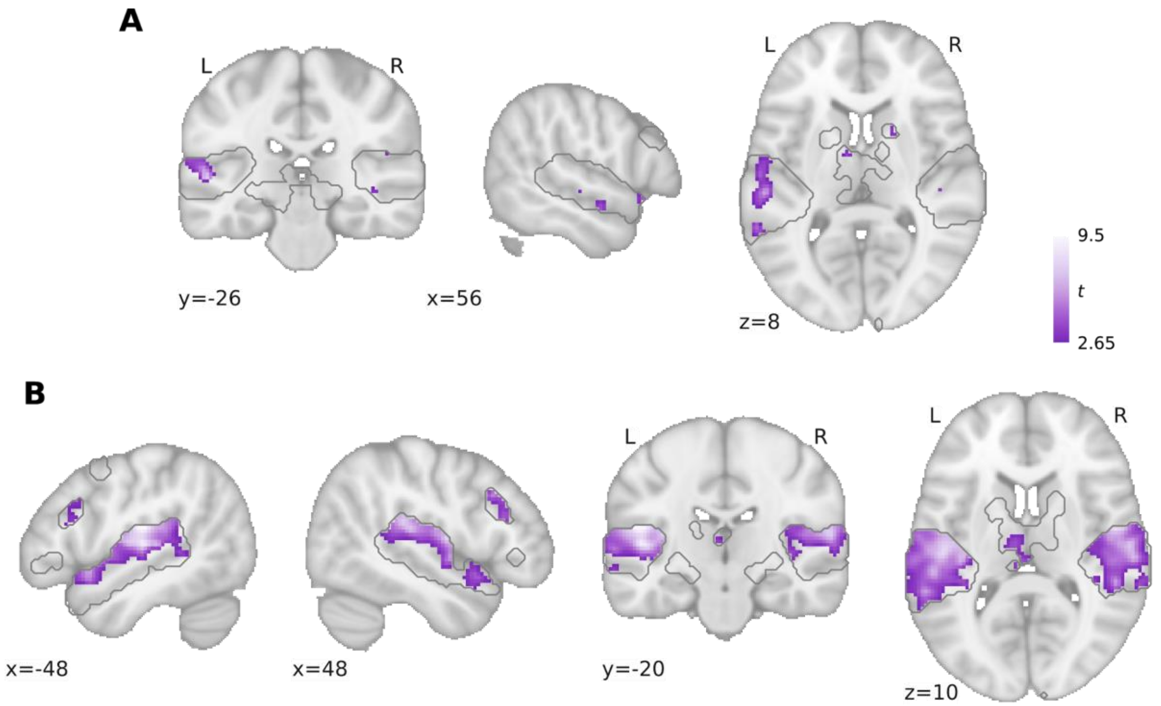


**Supplemental Figure 7. Effector decoding searchlight maps from Experiment 2. A.** Cluster-uncorrected searchlight map during the Delay epoch, which reveals no sub-threshold decoding clusters in right (ipsilateral) STG. This suggests that delay period decoding in Experiment 2 is contralateral in nature. **B.** Cluster-corrected searchlight map during the Execute epoch. Seven clusters were detected in the following regions: left STG (2751 voxels; peak x = -50, y = -20, z = 16, $t_{14}$ = 10.14), right STG (2063 voxels; peak x = -50, y = -30, z = 18, $t_{14}$ = 8.75), left thalamus (539 voxels; peak x = -6, y = -32, z = 0, $t_{14}$ = 16.32), cerebellum (527 voxels; peak x = 2, y = -82, z = -22, $t_{14}$ = 11.04), right inferior frontal gyrus (IFG; 202 voxels; peak x = 58, y = 20, z = 30, $t_{14}$ = 6.16), left IFG (191 voxels; peak x = -54, y = 18, z = 26, $t_{14}$ = 5.86), and left amygdala (152 voxels; peak x = 16, y = -14, z = -14, $t_{14}$ = 7.30). .

**Supplemental Table 1. ROI sizes used for analyses.**

| ROI | Size (2mm iso-voxels) |
|---|---|
| Left STG | 872 |
| Right STG | 832 |
| Left HG | 368 |
| Right HG | 318 |
| Left PT | 296 |
| Right PT | 281 |
| Left PMd | 96 |
| Right PMd | 110 |

#### *Behavioural Control Study for Experiment 1*

All Experiment 1 subjects participated in a behavioural testing session (performed outside the MRI scanner and before the experimental task) in which their eye fixations and forces corresponding to object grasping and lifting were measured as participants completed the experimental task. This testing session was used for participant screening and to determine, from an analysis of their object lifting and eye-movement behaviour, whether participants were, respectively, (1) maintaining in working memory the instructed hand information over the delay period of each event-related trial and, (2) able to reliably maintain fixation over the duration of an fMRI experimental run (thereby arguing against standard, alternative 'eye-movement confound' interpretations of the fMRI data). In this behavioural testing session, each participant completed 3 experimental runs, identical to those performed in the MRI scanner during the experimental testing session.

The experiment apparatus, equipment and setup was identical to that used in the MRI scanner, but testing was instead performed in the behavioural laboratory. Prior to beginning the behavioural experiment, participants received both verbal instructions and a demonstration by the experimenter as to how to correctly perform the object-directed actions. [Note that force measurements in this behavioural testing session were primarily taken only to provide additional confirmation that participants were capable of performing the task correctly.]

During this behavioural testing, an infrared video-based eye-tracking system (ETL 500 pupil/corneal tracking system, ISCAN Inc. Burlington, MA, USA), mounted below a headband, recorded the gaze position of the left eye at 240 Hz as the participant maintained gaze on the fixation LED. Gaze was calibrated using a two-step procedure: an initial 5-point calibration using ISCAN's Line-of-Sight Plane Intersection Software followed by a 25-point calibration routine. Twenty-five calibration points (4 mm-diameter circles) were shown on a cardboard frame, presented at the distance of the fixation point, and distributed over a region that incorporated the fixation point, the hand start location, and the location of the central object position. The ISCAN calibration converted raw gaze signals into pixels from the line-of-sight camera and the 25-point calibration converted pixels (i.e., the output of the ISCAN calibration) into the coordinates of hand workspace. Gaze was calibrated at the start of the experimental run and was checked following each block of trials so that, if necessary, gaze could be re-calibrated before starting a new test block.

### References


Huth, A. G., Nishimoto, S., Vu, A. T., & Gallant, J. L. (2012). A continuous semantic space describes the representation of thousands of object and action categories across the human brain. *Neuron*, *76*(6), 1210–1224.

Markiewicz, C. J., & Bohland, J. W. (2016). Mapping the cortical representation of speech sounds in a syllable repetition task. *NeuroImage*, *141*, 174–190.

Stelzer, J., Chen, Y., & Turner, R. (2013). Statistical inference and multiple testing correction in classification-based multi-voxel pattern analysis (MVPA): random permutations and cluster size control. *NeuroImage*, *65*, 69–82.
